# Vast heterogeneity in cytoplasmic diffusion rates revealed by nanorheology and Doppelgänger simulations

**DOI:** 10.1101/2022.05.11.491518

**Authors:** Rikki M. Garner, Arthur T. Molines, Julie A. Theriot, Fred Chang

**Affiliations:** Biophysics Program, Stanford University School of Medicine, Stanford, CA, USA; Department of Biology and Howard Hughes Medical Institute, University of Washington, Seattle, WA, USA; Department of Cell and Tissue Biology, University of California San Francisco, San Francisco, CA, USA; Marine Biological Laboratory, Woods Hole, MA 02543, USA

**Author notes:** Harvard Medical School, Department of Systems Biology, Boston MA 02115. Authors had equal contributions to the work (Rikki M. Garner, Arthur T. Molines). Authors had equal contributions to the work (Julie A. Theriot, Fred Chang). Corresponding authors (R.M.G.), (A.T.M.).

**Keywords:** Biological noise, diffusion, cytoplasmm viscosity, rheology, particle tracking, fission yeast *Schizosaccharomyces pombe*, simulations

## Abstract

The cytoplasm is a complex, crowded, actively-driven environment whose biophysical characteristics modulate critical cellular processes such as cytoskeletal dynamics, phase separation, and stem-cell fate. Little is known about the variance in these cytoplasmic properties. Here, we employed particle-tracking nanorheology on genetically encoded multimeric 40-nm nanoparticles (GEMs) to measure diffusion within the cytoplasm of the fission yeast *Schizosaccharomyces pombe*. We found that the apparent diffusion coefficients of individual GEM particles varied over a 400-fold range, while the differences in average particle diffusivity among individual cells spanned a 10-fold range. To determine the origin of this heterogeneity, we developed a Doppelgänger Simulation approach that uses stochastic simulations of GEM diffusion that replicate the experimental statistics on a particle-by-particle basis, such that each experimental track and cell had a one-to-one correspondence with their simulated counterpart. These simulations showed that the large intra- and inter-cellular variations in diffusivity could not be explained by experimental variability but could only be reproduced with stochastic models that assume a wide intra- and inter-cellular variation in cytoplasmic viscosity. The simulation combining intra- and inter-cellular variation in viscosity also predicted weak non-ergodicity in GEM diffusion, consistent with the experimental data. To probe the origin of this variation, we found that the variance in GEM diffusivity was largely independent of factors such as temperature, cytoskeletal effects, cell cycle stage and spatial locations, but was magnified by hyperosmotic shocks. Taken together, our results provide a striking demonstration that the cytoplasm is not “well-mixed” but represents a highly heterogeneous environment in which subcellular components at the 40-nm size-scale experience dramatically different effective viscosities within an individual cell, as well as in different cells in a genetically identical population. These findings carry significant implications for the origins and regulation of biological noise at cellular and subcellular levels.

**Significance:** Biophysical properties of the cytoplasm influence many cellular processes, from differentiation to cytoskeletal dynamics, yet little is known about how tightly cells control these properties. We developed a combined experimental and computational approach to analyze cytoplasmic heterogeneity through the lens of diffusion. We find that the apparent cytoplasmic viscosity varies tremendously – over 100-fold within any individual cell, and over 10-fold among individual cells when comparing averages of all particles measured for each cell. The variance was largely independent of temperature, the cytoskeleton, cell cycle stage, and localization, but was magnified under hyperosmotic shock. This suggests that cytoplasmic heterogeneity contributes substantially to biological variability within and between cells, and has significant implications for any cellular process that depends on diffusion.

**Graphical abstract:** 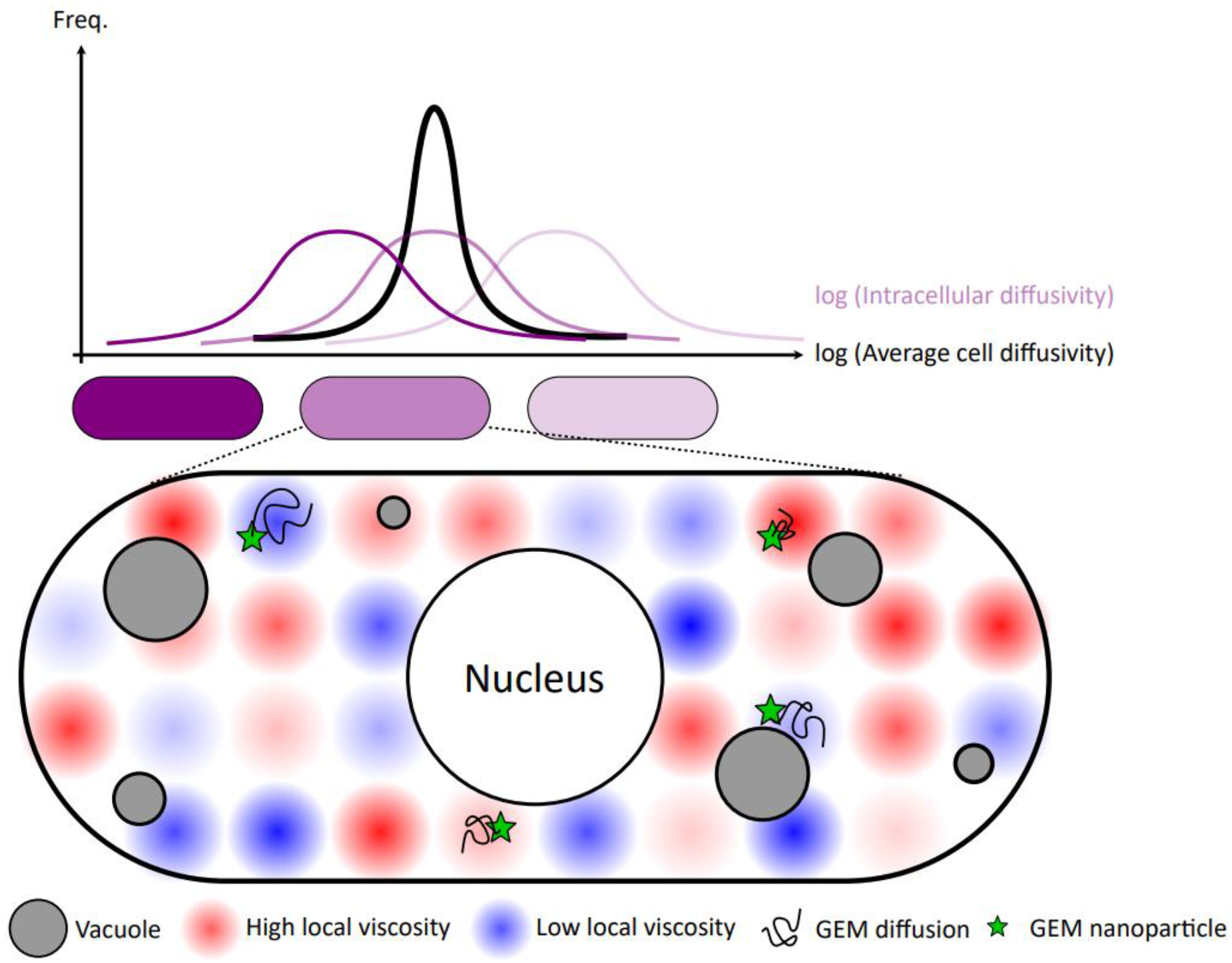

## Introduction

Life at the molecular scale is stochastic, with macromolecules continually being jostled by Brownian motion. This emergence of “biological noise” at the molecular level permeates all aspects of cell biology, inducing stochastic fluctuations in subcellular processes and driving natural variation among cells in a population. Previous work has outlined critical roles for biological noise in signaling (1), cell size control (2–4), organelle size scaling (5–7), and gene expression (8–11). In general, biological noise presents a challenge to cellular homeostasis and signaling mechanisms, and is often suppressed in order for biological functions to be robust. For example, signaling frequently depends on strong amplification of initially weak signals, which can erroneously amplify noise unless proofreading mechanisms are in place (1). However, biological noise can also confer a selective advantage. In a fluctuating and unpredictable environment, biological variation between cells in an isogenic population can ensure population-level survival (12, 13).

One potentially significant source of biological noise that has been largely ignored is that of heterogeneity in the cell cytoplasm. The cytoplasm is composed of a highly diverse and actively-mixed assembly of resident macromolecules of various size (14, 15), charge (15), and hydrophobicity (16). The complexity of the cytoplasmic milieu could influence molecules’ behavior locally. Indeed, spatiotemporal heterogeneity in the diffusion of particles has been observed in multiple contexts such as *E. coli*, fungi, mammalian cells and even *Xenopus* egg extract using methods ranging from fluorescence correlation spectroscopy (FCS) to particle tracking (17–31).

Heterogeneity of cytoplasmic properties have potentially far-reaching effects in cell biology, as the cytoplasm hosts a wide variety of critical molecular processes ranging from protein synthesis and turnover, to cytoskeletal transport and force production, metabolism, and beyond (32, 33). Further, changes to physical cytoplasmic properties such as the macromolecular density, viscosity, and degree of crowding have been shown to impart widespread effects within the cell—including sudden and significant impacts on growth and viability (34, 35). For example, altering cytoplasmic crowding by changing the concentration of ribosomes has strong effects on phase separation (36), and high osmotic shocks can completely halt microtubule dynamics (37). Additionally, alterations in cytoplasmic density have been implicated in cellular aging and senescence (38) and differentiation (39).

Here we establish a combined experimental and computational approach to examine cytoplasmic heterogeneity through the lens of diffusion. Single particle motion-tracking allows for a robust quantification and statistical analysis of particle behavior, revealing variations between particles which would otherwise be averaged out in bulk measurements obtained in photobleaching (e.g. FRAP) and FCS (33, 40) experiments. Further, this kind of “passive” rheology approach requires minimal perturbations to the cell.

Previously, particle tracking rheology on fluorescent proteins has proven difficult due to their fast diffusion rates and tendency to photobleach. The development of GEMs (Genetically Encoded Multimeric nanoparticles (36)), has enabled large improvements on this front. These bright and photostable protein spheres are expressed as fluorescently-tagged monomers which self-assemble into hollow shells of nearly-uniform size and shape (36, 41). Because each particle contains tens of fluorescent proteins, they can be tracked for relatively long periods of time without photobleaching. Additionally, the near-diffusive movements of GEMs suggest they do not interact strongly with eukaryotic cellular components, making them ideal reagents for rheological studies (22, 36). Critically, their relatively large size and slow diffusion rates - comparable to large protein complexes such as ribosomes - allow GEMs to be tracked using modern high-speed cameras (which is still not attainable for individual fluorescent proteins). Initial studies have established their utility in quantitatively probing diffusion and crowding in the cytoplasm and nucleoplasm in various cell types including yeast and mammalian cells (22, 36, 41–45).

The fission yeast *Schizosaccharomyces pombe* provides an excellent model system for the study of cytoplasmic heterogeneity because of their uniformity in many other aspects of their cell biology. In standard laboratory conditions, these rod-shaped cells exhibit very tight distributions in their cell size at division (CV ~ 6% (3, 4)) and cell shape (46–48), as well as cell cycle progression and intracellular density (CV ~ 10% (49)). The relatively low phenotypic variability within and between fission yeast cells permits the study of cytoplasmic heterogeneity in a well-controlled system in the presence of minimal confounding factors.

Using live cell high-speed imaging and quantitative tracking of 40 nm-diameter GEMs in *Schizosaccharomyces pombe*, we measured cytoplasmic diffusivity for thousands of individual particles. These data revealed large heterogeneity in diffusion coefficients both within single cells as well as between cells in the population. To analyze this variability, we developed an automated pipeline, which we call the Doppelgänger Simulation approach, to reproduce our experimental results computationally using simulations of diffusion, and assay heterogeneity using statistical techniques for analysis of variance. Using these methods, we showed that orders of magnitude of variability in GEM cytoplasmic diffusivity within and between cells arose from an equally wide distribution of cytoplasmic viscosity. This variance was not affected by temperature, the cytoskeleton, or cell size, but was increased by hyperosmotic shock. Our studies support a growing body of evidence that the cytoplasm is not physically well-mixed (17–20, 22, 23, 50) and reveal this heterogeneity in diffusion as an important potential source of biological noise.

## Results

### Statistical characterization of cytoplasmic GEM particle diffusion in fission yeast

To assay cytoplasmic diffusion in fission yeast, we expressed 40 nm-diameter GEM nanoparticles in wildtype *S. pombe* from a multicopy plasmid on an inducible promoter (37). Tuning the expression of the GEMs construct allowed us to titrate particle formation to a small number of particles (<10) per cell. To reduce environmental variability, these cells were grown at 30 °C under optimal conditions in shaking liquid cultures to exponential growth phase and mounted in imaging chambers with fixed dimensions under constant temperature and imaged acutely. Using variable angle epifluorescence microscopy (VAEM) (36, 37, 44, 51) we tracked GEM particle motion at 100 Hz for 10 s, as described previously (36, 37, 44) (Fig. 1a-b, *Methods*). Each field of view (FOV) contained multiple cells that were individualized post-acquisition. Images were manually curated to eliminate from the data set a small subset of cells that had died, exhibited grossly abnormal morphologies, or contained a single bright aggregate of GEM particles. From a dataset of 145 cells, 3681 tracks were analyzed, with an average of 25 ± 10 (AVG ± SD) tracks per cell, a mean step size of 104 ± 72 nm (AVG ± SD), and a mean track length of 273 ± 268 ms (AVG ± SD) (Fig. 1c).

From these trajectories we computed the time-averaged, ensemble-averaged (i.e., track-averaged) mean squared displacement (MSD) as a function of time interval, and fitted the resulting MSD curve to a power law (Fig. 1d, *Methods*). MSD analysis showed that GEM particle motion in the cytoplasm was largely diffusive (MSD ≈ Dτ), following a robust power law with apparent diffusivity D_app,100ms_ 0.3 ± 0.01 μm^2^/sec (AVG ± 95% CI) and anomalous diffusion exponent α = 0.92 ± 0.02 (AVG ± 95% CI), which were similar to previously published measurements in fission yeast (43, 44). The diffusivity of the 40 nm GEMs in the cytoplasm was roughly 40 times slower than the theoretical prediction for simple Stokes-Einstein diffusion in water -- and corresponded to the particle’s expected diffusion rate in a 75% glycerol solution in water. We note that diffusion along the long and short axes of the cell were comparable by our measurements (Supp. Fig. 1a), and we found that the MSD plots do not plateau, indicating that diffusion of the GEMs was not confined on timescales less than a second (e.g., most particles do not run into the cell wall within the measured time window).The time-averaged, ensemble-averaged (i.e., track-averaged) velocity autocorrelation of particle trajectories was also consistent with simple unconstrained diffusion (Fig. 1e). Notably, the autocorrelation plot lacked the characteristic negative peak associated with subdiffusive motion and viscoelastic response seen in other systems (Supp. Fig. 1b-c) (52–55). Therefore, at least with this approach at this 40-nm size scale, we detected no elastic response in the yeast cytoplasm.

**Figure 1:**
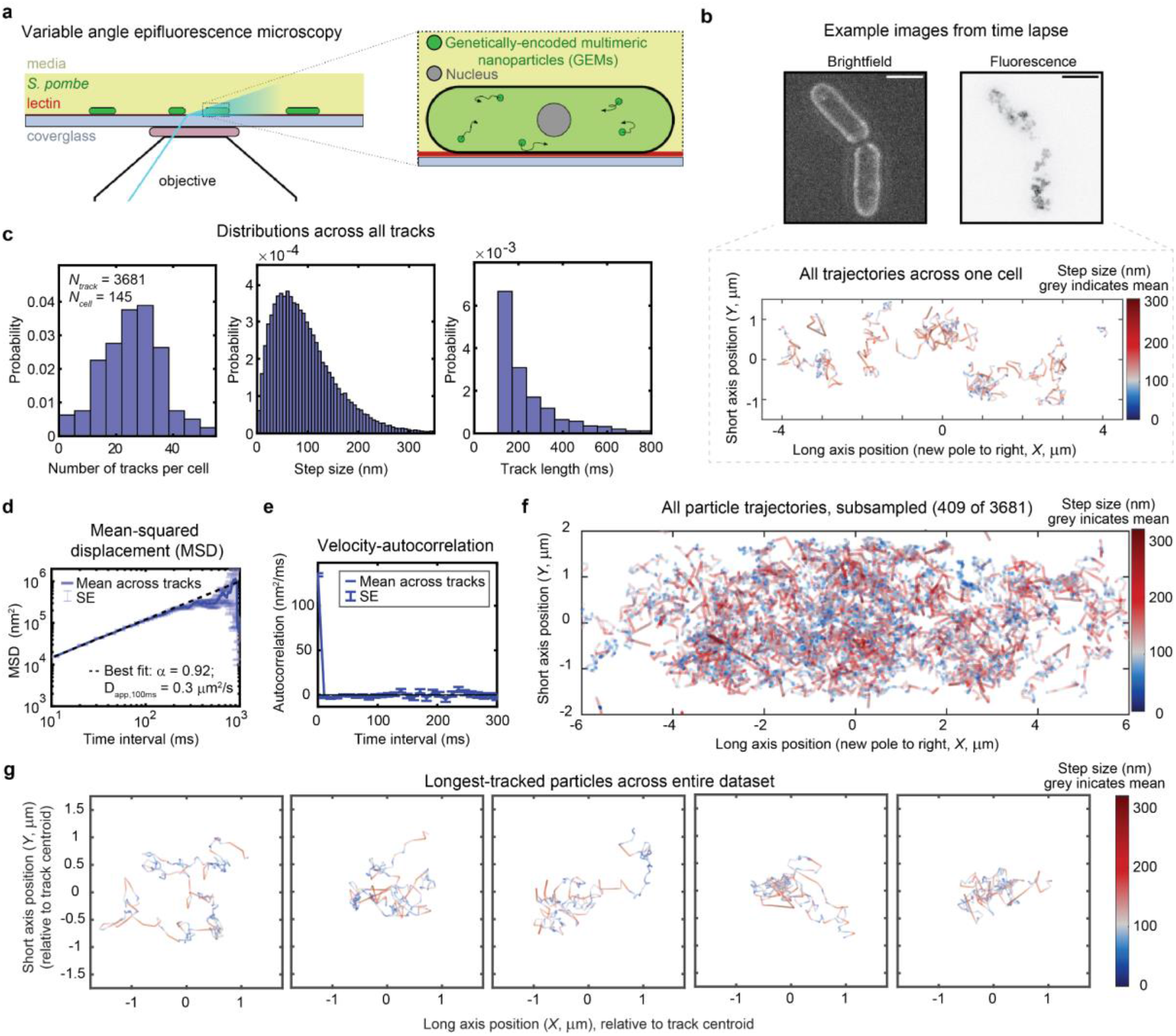
High-speed particle-tracking nanorheology of GEMs allows detailed statistical analysis of cytoplasmic diffusion. (**a**) Schematic of the experimental imaging set-up. (**b**) Example brightfield image (top left) and maximum intensity projection through time of the GEM particle fluorescence (top right) for one representative field of view, alongside the measured nanoparticle trajectories (bottom) for the upper cell in the image. Trajectories are colored by the step size of the particle in nanometers between each time frame of the movie. Gray indicates the mean step size across all tracks in the dataset. Scale bar is 5 μm. (**c**) Histograms of the number of tracks per cell (left), the step-size for all time-points (middle), and the duration of time that each particle was tracked (right). Note that tracks shorter than 10 time-points were not included in the analysis. (**d**) The mean squared displacement (MSD) of the particle tracks. The time-averaged MSD was first calculated individually for each track, and then a second averaging was performed to find the (ensemble averaged) MSD across all tracks. Note the logarithmic scale along the x- and y-axes. (**e**) The average velocity autocorrelation across all article tracks. Averaging was performed in the same order as the MSD. (**d-e**) Error bars represent the standard error. (**f**) Plots of particle trajectories drawn from many experiments and cells, randomly subsampled for better visibility of individual particle behaviors. Subsampled trajectories include at least one track from 141 of the 145 cells in the dataset. Gray indicates the mean step size across all tracks in the dataset. (**c-f**) Dataset includes 3681 tracks among 145 cells, recorded from 5 different samples and over 3 different days. (**g**) Individual trajectory plots for five of the longest-tracked particles (in time), excluding stationary particles. Color scaling of the step size was identical in all panels included in **f-g** (using the mean and standard deviation of the step size across the entire dataset).

### Cytoplasmic diffusivity spans orders of magnitude

We next analyzed individual particle tracks, which revealed a rich phenotypic variability (Fig. 1f-g) that was obscured by the ensemble averaging-based analysis described above (e.g., MSD - Fig. 1d). Notably, even within a single trace, individual particles exhibited large fluctuations in their step size (Fig. 1g). To investigate the variety of comportment displayed by individual particles, we calculated and fit the time-averaged MSD individually for each track (Fig. 2a) and the time-averaged, ensemble-averaged MSD over all tracks in each cell (Fig. 2b). These data showed that variability in particle motion ranged over orders of magnitude; fits of particle and cell MSDs (Fig. 2c-d) showed that diffusivity follows a long-tailed, *log-scale* distribution, consistent with Brownian motion in a heterogeneous environment (56, 57). The distribution of apparent diffusivities exhibited a single peak (Fig. 2c-f), which appeared more normally-distributed on a log scale (Fig. 2f) than on a linear scale (Fig. 2e). Therefore, we performed all further statistics and visualization on the log10 of the fitted apparent diffusivities. The median of the diffusivity distribution in log space, which we then converted to linear space (see Methods), corresponded to a diffusivity of 0.29 μm^2^/sec for the track-wise distribution and 0.33 μm^2^/sec for the cell-wise distribution, both similar to the bulk estimate. The standard deviation of the diffusivity distribution in log space (representing the number of orders of magnitude spanned by the dataset) can be converted to linear space as a fold-range at 2.5 standard deviations away from the median (see Methods), giving a 392-fold range across tracks and 11-fold range across cells. We chose 2.5σ as our cutoff as it gave a range consistent with our outlier estimation algorithm (Fig. 2c, see caption). Overall, we showed that diffusivities vary by over 2.5 orders of magnitude among individual GEMs and one order of magnitude among cells. Hereafter we use the terms *inter*cellular variation to refer to the spread of the cell-wise diffusivity (Fig. 2c, right) and *intra*cellular variation to indicate the spread in the track-wise diffusivity (Fig. 2c, left).

**Figure 2:**
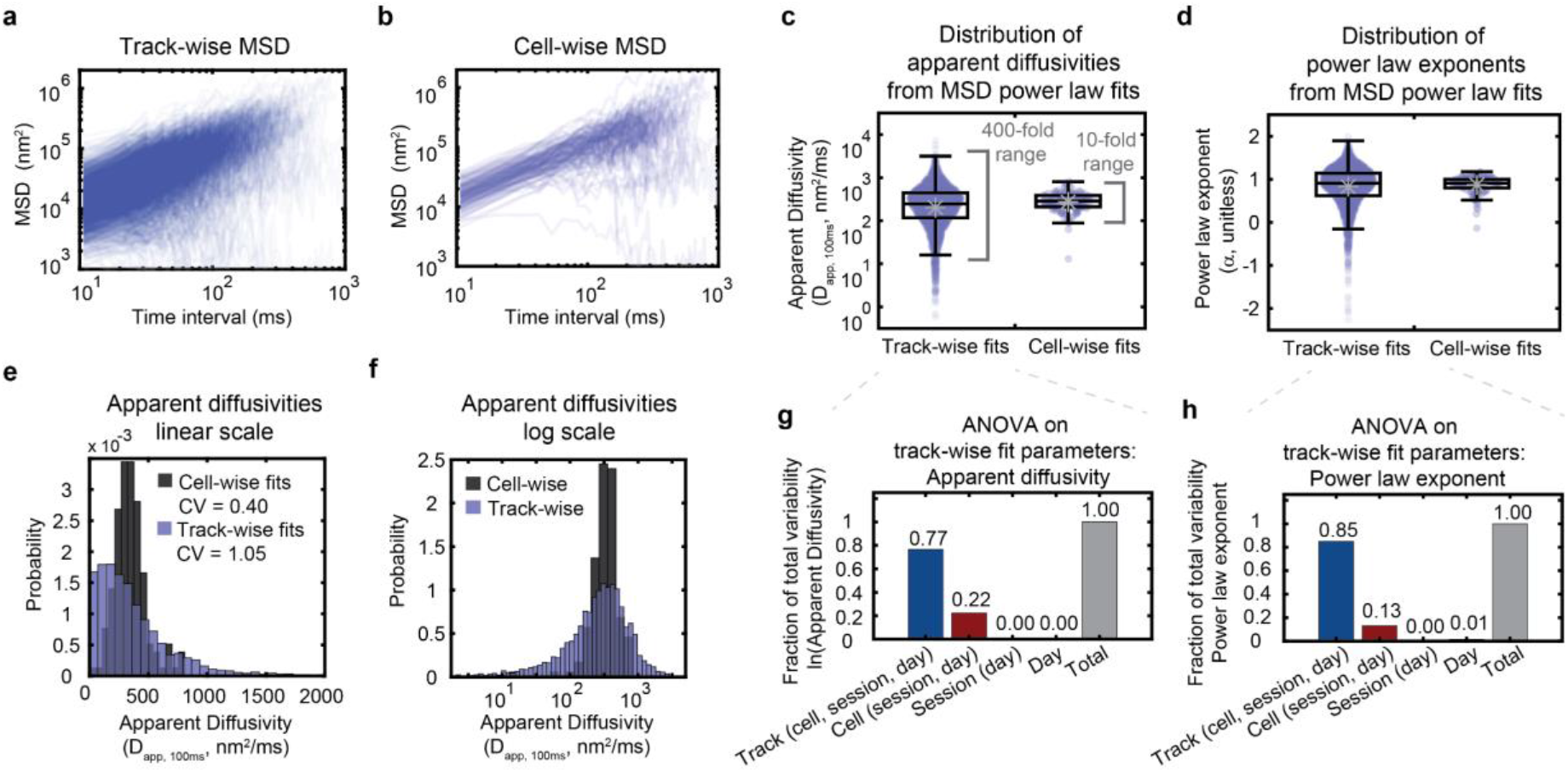
GEM diffusivity varies over 400-fold across tracks and 10-fold across cells. (**a-b**) Mean squared displacements averaged either (**a**) by track (averaged over time for each track), or (**b**) by cell (averaged over time for each track and then averaged across all tracks in each cell). Note the logarithmic scale along the x-and y-axes. (**c-d**) Apparent diffusivities (**c**) and power law exponents (**d**) calculated from fits of the track-wise and cell-wise MSDs to a power law. Note the logarithmic scale along the y-axis. Boxplots: Central line, median; grey dot, mean; boxes, 25th and 75th percentiles; whiskers, furthest data point that is not an outlier; outliers, any point that is more than 1.5 times the interquartile-range past the 25th and 75th percentiles. (**e-f**) The same distributions of the fitted apparent diffusivities plotted in (**c**), now plotted as a histogram either on a linear scale (**e**) or on a log scale (**f**). Probabilities represent the probability density per histogram bin width, such that the sum of the bin heights multiplied by the bin width equals 1. (**g-h**) Results from a nested ANOVA performed on track-wise fits of diffusivities (**g**) and power law exponents (**h**). The amount of the experimentally-observed variance that can be explained by track-to-track, cell-to-cell, imaging session-to-session, and day-to-day variability is plotted as a fraction of the total variance. (**a-h**) Dataset is identical to that shown in Fig. 1 **c-f**, including 3681 tracks among 145 cells, recorded from 5 different samples and over 3 different days.

To understand whether variation arose from cell-to-cell variability, from different microenvironments within a single cell, or from experimental day-to-day variation, we performed an Analysis of Variance (ANOVA) on the track-wise diffusivity measurements (Fig. 2g). The ANOVA revealed that the vast majority (~80%) of the measured spread in diffusivity came from intracellular variation (i.e., *Track* in Fig. 2g), but there was also a significant amount of variance (~20%) explained by cell-to-cell variability (i.e., *Cell* in Fig. 2g). Only < 1% could be attributed to experiment-to-experiment variability. Similar results were observed for the fitted anomalous diffusion exponent (Fig. 2d, h), which was not surprising given the strong correlation between the fitted apparent diffusivities and power law exponents in our dataset (Supp. Fig. 2a).

Another common way to differentiate sources of noise in biological data is to separate the observed spread into intrinsic (uncorrelated within cells) and extrinsic (correlated within cells) components (9, 58). Our ANOVA results suggested that noise in this system was almost entirely intrinsic, as ~80% of the variation was maintained after controlling for cell-to-cell variability. Indeed, by plotting the apparent diffusivities of random pairs of GEM particles, where each pair is randomly chosen from particles within a single cell, we found that the noise had only a very weak correlation between particles within the same cell (Spearman correlation: r = 0.21, p = 5*10^-35^, Supp. Fig. 2b). We noted that the large intercellular and intracellular variation observed in our data cannot be explained by differences in GEM particle expression levels, as the mean apparent diffusivity among track-wise diffusivity fits within a cell was not significantly correlated with the number of tracks in the cell (Spearman correlation: r = −0.003, p = 0.97, Supp. Fig. 2c), and the coefficient of variation among trackwise diffusivity values within a cell was only very weakly correlated with the number of tracks in the cell (Spearman correlation: r = 0.25, p = 0.004, Supp. Fig. 2d). In addition, the coefficient of variation of particle diffusivities within each cell was ~1, and was uncorrelated with the mean particle diffusivity across all particles in the cell, consistent with Poisson statistics (Spearman correlation: r = −0.1, p = 0.15, Supp. Fig. 2e).

### Stochastic simulations demonstrate that spread is not due solely to statistical measurement noise

As diffusion is an inherently stochastic process, we next explored whether the measured variation in particle mobility was due to statistical properties of our measurements. It is known, for example, that datasets with shorter track lengths will produce wider distributions of measured diffusivities (55). We therefore developed what we called the Doppelgänger Simulation (DS) approach, employing a custom algorithm to automatically read in and replicate the experimental measurement statistics *in silico* cell-for-cell and track-for-track (Fig. 3a). With DS, simulated cells have the exact same cell length and number of tracks as their experimentally-measured counterparts, and each simulated track has an identical length (i.e., number of time points tracked) to the associated experimental trajectory. This straightforward and powerful approach allowed us to produce simulated data that could be directly compared to the experimental tracks and analyzed using identical statistical methods.

**Figure 3:**
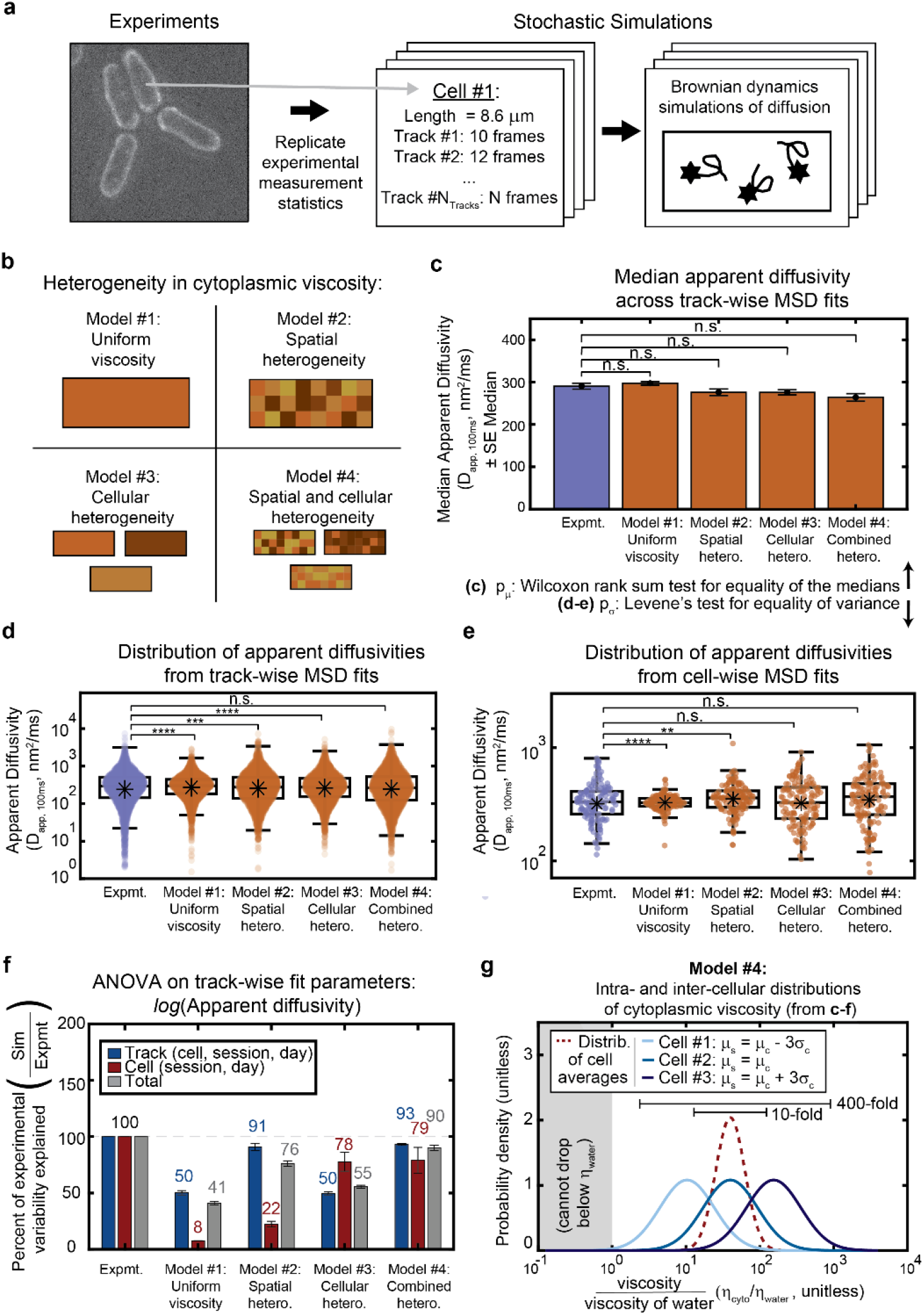
Stochastic simulations reveal both spatial and cellular heterogeneity in viscosity are required to reproduce experimentally observed variation. (**a**) Schematic of the Mirror Image Simulation approach. Each experimentally-measured cell and particle were reproduced one-to-one in the simulated dataset, with every simulated cell having the same long-axis length as its experimentally-measured counterpart, and each particle being tracked for the same amount of time. (**b**) Schematic demonstrating different types of heterogeneity in cytoplasmic viscosity included in each of the four models. Note: the choice of physical domain size for spatial heterogeneity in (c-g) is 1000 nm. (**c**) Median apparent diffusivity (averaged across all tracks) plotted for the experimental dataset as well as each model. Error bars represent the standard error of the median. Significance stars represent the result of the Wilcoxon rank sum test for equality of the medians. (**d-e**) Distributions of apparent diffusivities calculated from fits of the track-wise (**d**) or cell-wise (**e**) MSD curves displayed for the experimental data as well as each of the models. Note the logarithmic scale along the y-axis. Boxplots are drawn as in Figure 2. Significance stars represent the result of Levene’s test for equality of variance. (**c-e**) * p<0.05. **p<0.01, *** p<0.001, **** p<0.0001. (**f**) Results from a nested ANOVA performed on track-wise fits of diffusivities (**d**). The percent of the experimentally-observed cell-to-cell and track-to-track variability that can be explained by each of the models. (**g**) The distribution of cytoplasmic viscosities, shown relative to the viscosity of water, needed to most closely reproduce the experimental data (i.e., simulations from Model #4, using the same parameters used to generate (**c-f**)). Histograms are shown for the distribution of average cell viscosities (intercellular heterogeneity, red dashed line) and the distribution of intracellular viscosities for three example cells (blue lines of varying darkness). The examples include a cell whose average viscosity equals that of the cell-wide average (medium blue line), a cell with an average viscosity three standard deviations above the cell-wide average (dark blue line), and a cell with an average viscosity three standard deviations below the cell-wide average (light blue line). Note the logarithmic scale along the x-axis. The simulation did not allow viscosities below that of water.

Using DS, particle motion was then recapitulated using stochastic Brownian dynamics simulations of diffusion inside a box representing the exterior cell boundary (Fig. 3, Tables 2–3). We opted for a simple diffusion model because GEM particle motion is observed experimentally to be largely diffusive (Fig. 1d) and did not display characteristics of constrained or viscoelastic behavior (Fig. 1d-e). The model assumes an average cytoplasm viscosity forty times that of water, giving a mean diffusivity of 0.35 μm^2^/s that closely matches that of the experimental data. The simplest iteration of the model (Fig. 3b-f, Model #1: uniform viscosity), which we will hereafter refer to as the uniform viscosity model (due to its assumption of constant viscosity within and among cells), accounted for only a fraction of the experimentally-measured spread in GEM particle mobility (Fig. 3d-f) – including ~50% of the track-to-track variability in diffusivity, and <10% of the cell-to-cell variability, as measured by ANOVA (Fig. 3f). We therefore concluded that neither the stochastic nature of diffusion nor the statistical properties of our experimental measurement statistics were the major source of heterogeneity in GEM particle diffusion.

**Table 1:**
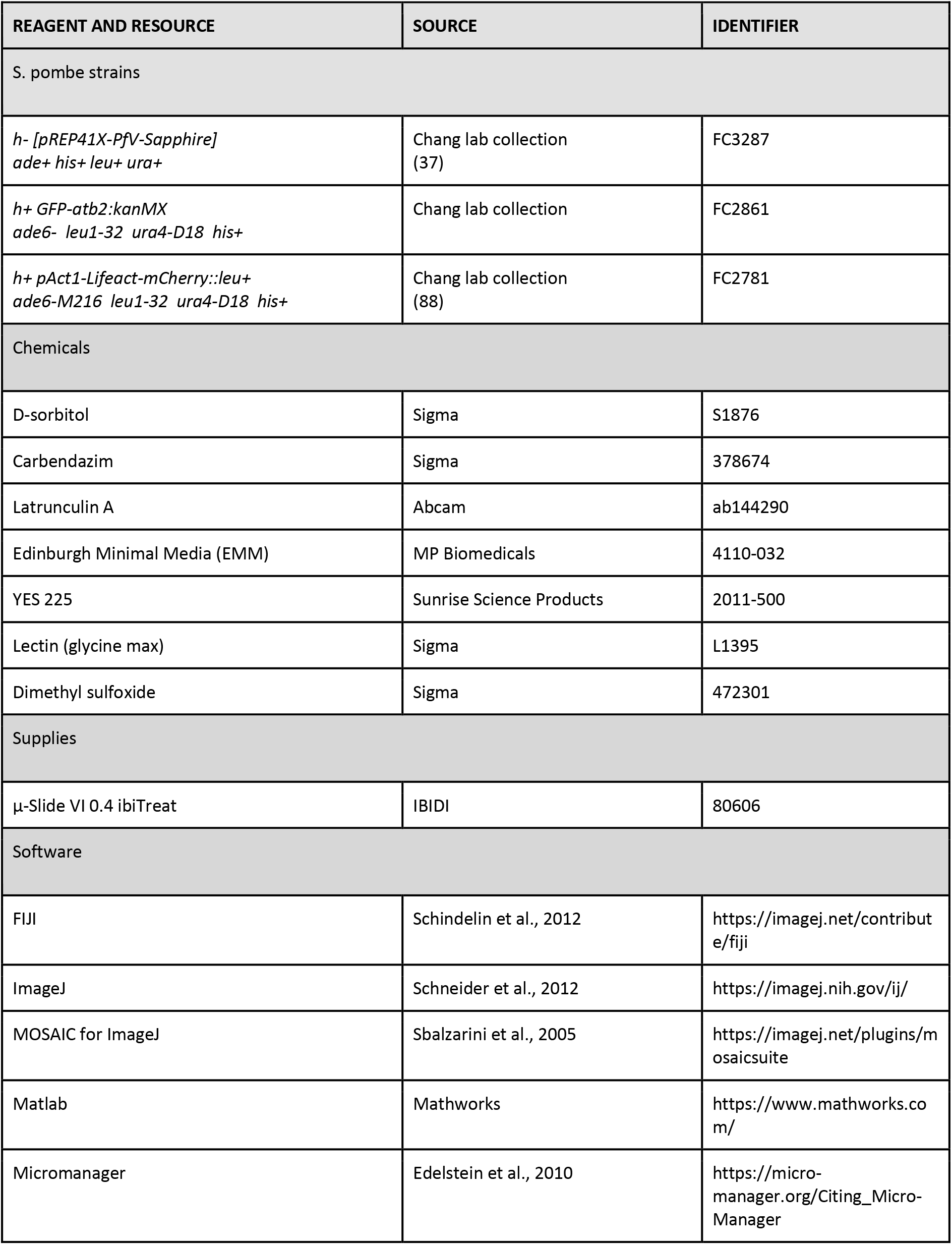
Reagents and Resources.

**Table 2:**
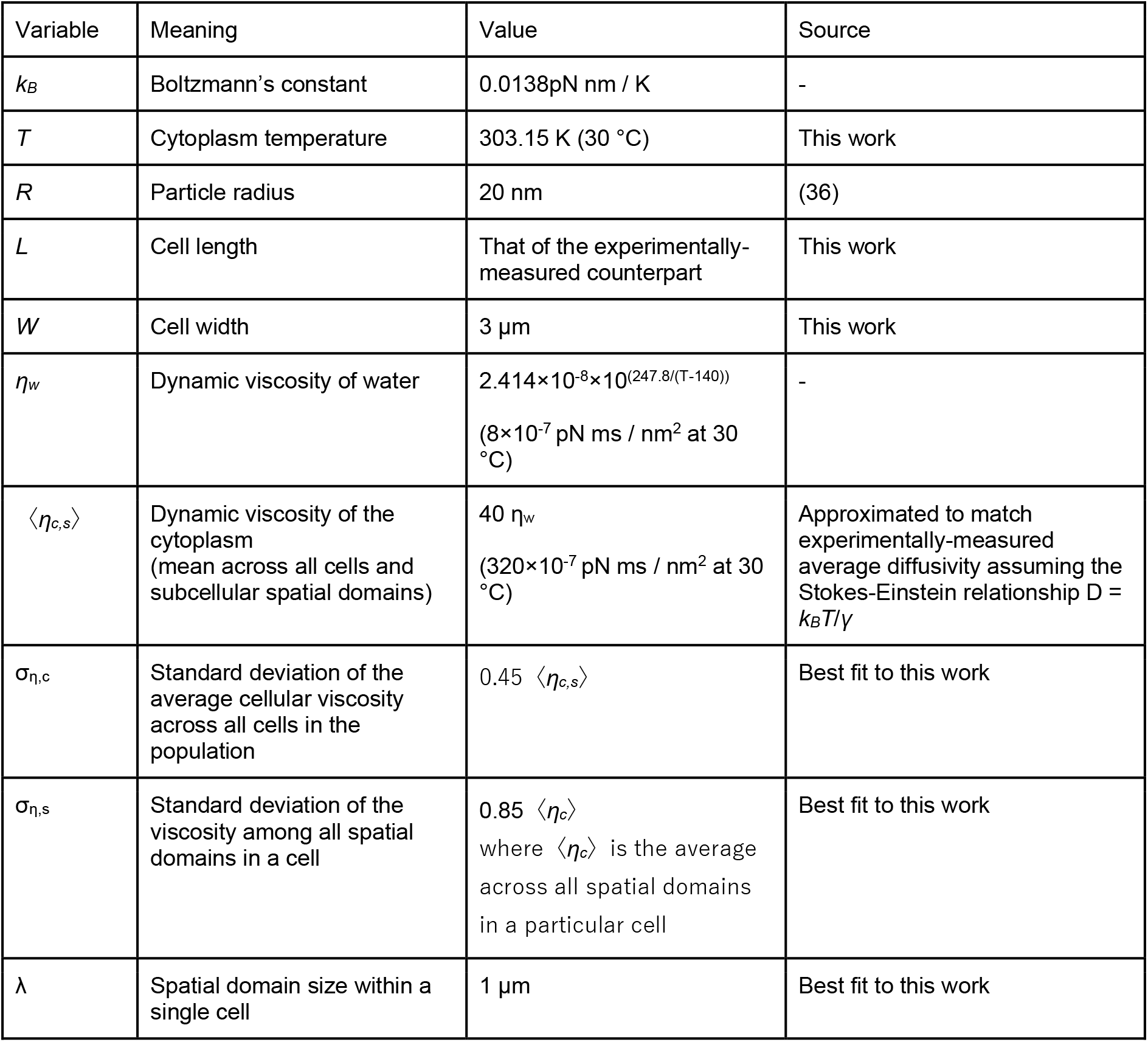
Model Parameters - Input parameters.

**Table 3:** Model Parameters - Derived parameters.

| Variable | Meaning | Value |
| --- | --- | --- |
| $k_B T$ | Thermal energy | 4.18 pN nm |
| $\langle \gamma \rangle$ | Viscous drag coefficient given Stokes' law<br>(for the average particle)<br>$\gamma = 6\pi \langle \eta_c \rangle R$ | 0.012 pN ms / nm (at 30 °C) |
| $\langle D_{c,s} \rangle$ | Diffusivity<br>(averaged across all cells and spatial domains)<br>$D = k_B T / \gamma$ | 350 nm<sup>2</sup>/ms |

### Simulations suggest that heterogeneity in diffusion must arise from an equally vast spread in cytoplasmic viscosity

As the data set of experimentally-measured GEM particle motion fitted well to a model of simple diffusion, there were only a finite number of sources in this simple model from which heterogeneity in mobility could arise. The major parameter defining diffusion is the diffusivity, *D*, which theoretically (by the Stokes-Einstein equation) is simply equal to the ratio of the thermal energy, *k_B_T*, to the viscous drag on the particle, *γ*. For a spherical particle, *γ = 6πηR*, where *η* is the viscosity of the cytoplasm and *R* is the radius of the particle (59). Of these parameters, viscosity is the only parameter that could be varying within and between cells, as temperature is held constant and the radii of GEM nanoparticles have been shown by electron microscopy to be fairly uniform (CV ≈ 0.1) when expressed in mammalian and budding yeast cells (36).

We therefore generated three other versions of our model incorporating viscosity variation (Fig. 3b), while keeping the mean viscosity (and thus mean particle diffusivity) constant (Fig. 3c). In one version, which we refer to as the spatial heterogeneity model (Fig. 3, Model #2: spatial heterogeneity), we aimed to explore whether intracellular spatial variations in viscosity could account for the experimentally measured spread in diffusivity. A spatially varying viscosity was consistent with the observation that individual particles can display significant variations in step size within a single track (Fig. 1g). This model assumed that viscosity varies across the cell with a fixed domain size, approximated by a grid of discrete viscosity domains where each region was randomly assigned a distinct viscosity value. The average cellular viscosity was held constant. In another variation of the model, termed the cellular heterogeneity model, viscosity was uniform within each cell, but the uniform viscosity value varied between cells (Fig. 3, Model #3: cellular heterogeneity). Finally, we developed a fourth model combining both intracellular and intercellular heterogeneity, which we called the combined heterogeneity model (Fig. 3, Model #4: combined heterogeneity). In all three variations on the original model, viscosity values were chosen from a log-normal distribution, mimicking the distribution of the experimentally-measured step sizes (Fig. 1c, middle) and diffusivities (Fig. 2f).

We then ran each model multiple times to account for their stochastic nature and assayed whether each model could reproduce the experimentally-observed spread in diffusivity (1) as measured by ANOVA (Fig. 3f), and (2) such that the variance was not statistically significantly different from the experiments according to Levene’s test for equality of variances (Fig. 3d-e)(60). While the spatial heterogeneity model could only account for the track-to-track variation in experimentally-measured diffusivity (but not the cell-to-cell variation), and the cellular heterogeneity model could only reproduce the cell-to-cell variation (but not the track-to-track variation), only the model combining both spatial and cellular heterogeneity could fully reproduce the amount of spread observed in the experimental data (Fig. 3d-f). Further, only a viscosity variation spanning orders of magnitude (Fig. 3g) could quantitatively recapitulate the experimentally-measured spread. In particular, the viscosity was required to vary 10-fold (on average) among cells, 100-fold within any individual cell, and 400-fold across the dataset in order to best match the experiments. Importantly, this extreme degree of heterogeneity in viscosity was required regardless of the choice of characteristic length scale for the spatial variation (Table 4, Supp. Fig. 3). Overall, our simulations showed that our data is best explained by a model in which the effective viscosity experienced by cytoplasmic GEM particles varies drastically within and between cells.

**Table 4:** Model Parameters - Best fit parameters for each spatial heterogeneity domain size in order to match the experimentally-observed mean and variance in GEM diffusivity.

| Spatial domain size within a single cell $\lambda$ | Mean cytoplasm viscosity $\langle \eta_{c,s} \rangle$ | Standard deviation of the average cellular viscosity across all cells in the population $\sigma_{\eta,c}$ | Standard deviation of the viscosity among all spatial domains in a cell $\sigma_{\eta,s}$ |
| --- | --- | --- | --- |
| 100 nm | 50 $\eta_w$ | 0.375 $\langle \eta_{c,s} \rangle$ | 1.1 $\langle \eta_c \rangle$ |
| 300 nm | 39 $\eta_w$ | 0.4 $\langle \eta_{c,s} \rangle$ | 1.0 $\langle \eta_c \rangle$ |
| 600 nm | 38 $\eta_w$ | 0.4 $\langle \eta_{c,s} \rangle$ | 0.9 $\langle \eta_c \rangle$ |
| 1000 nm | 40.5 $\eta_w$ | 0.4 $\langle \eta_{c,s} \rangle$ | 0.8 $\langle \eta_c \rangle$ |
| 3000 nm | 41.5 $\eta_w$ | 0 $\langle \eta_{c,s} \rangle$ | 0.775 $\langle \eta_c \rangle$ |

### Spatial heterogeneity in cytoplasm viscosity can lead to ergodicity breaking

Heterogeneity in diffusion is frequently observed in non-ergodic systems, due to the fact that individual particles within the system exhibit distinct behaviors compared to the ensemble average (27–29, 61, 62). The experimentally-observed spread in GEM diffusivity thus suggested that the cytoplasm may represent a non-ergodic system. To test this hypothesis, we assayed the ergodicity of the GEM diffusion. A hallmark of a non-ergodic system is that the ensemble-averaged (EA) MSD diverges from time-ensemble-averaged (TEA) MSD (62). In comparing the EA MSD with the TEA MSD of our experimental dataset (Fig. 4) (62), we found that GEM diffusion was indeed weakly non-ergodic (Fig. 4a). In particular, the EA and TEA MSD exhibited a ~40% difference at short times, which decreased to ~10% at longer times (Fig. 4d).

To determine the origin of the weak non-ergodicity of GEM particle diffusion, we returned to our Doppelgänger simulations. As expected, we found that simulations with no heterogeneity in viscosity resulted in perfectly ergodic diffusion (Fig. 4b,e). However, simulations with both spatial and cellular heterogeneity (Model #4) were able to reproduce the experimentally-observed non-ergodicity (Fig. 4c), including quantitative features of the decay in non-ergodicity at long times (Fig. 4f).

**Figure 4:**
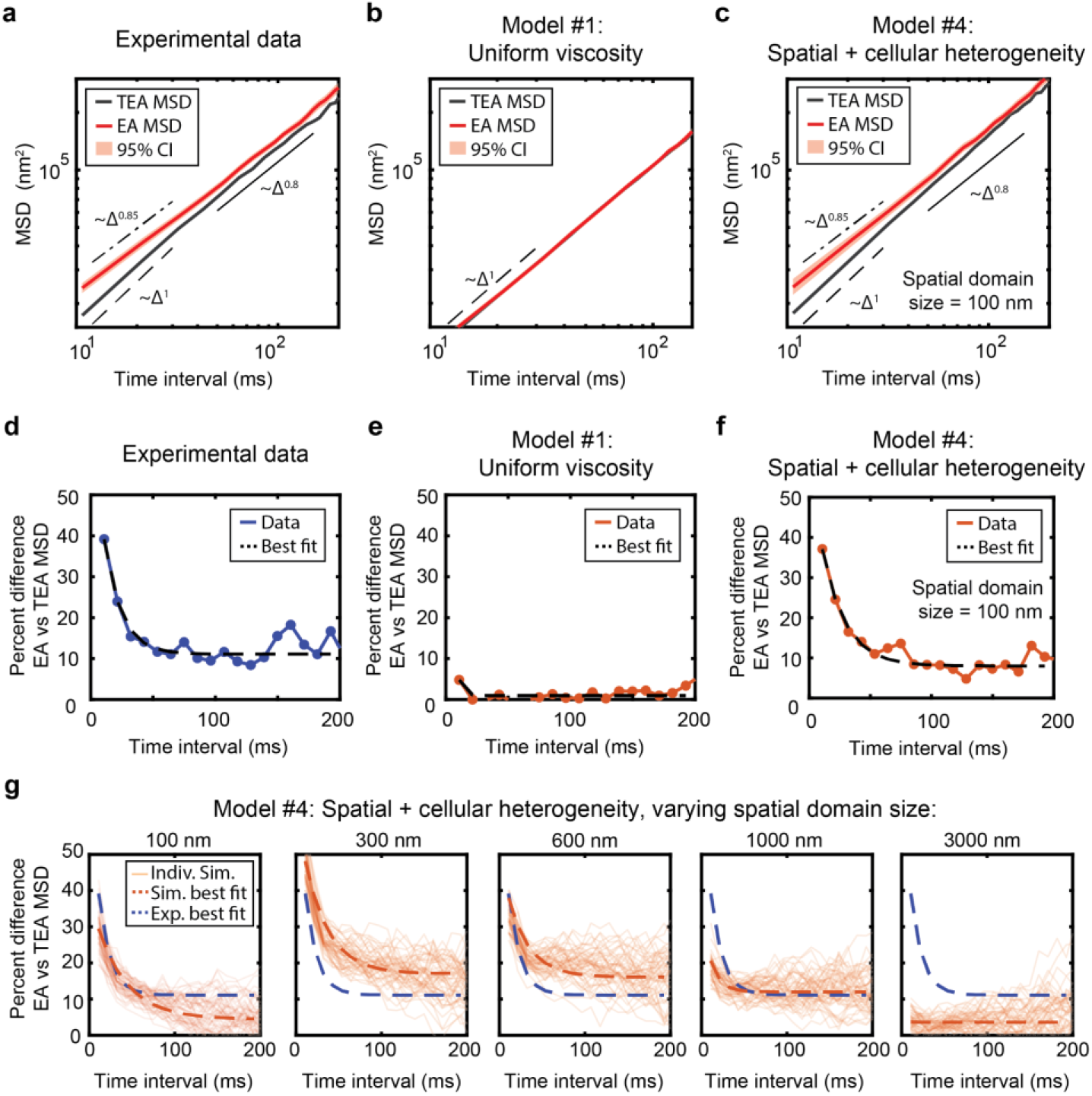
Weak non-ergodicity of GEM diffusion can be explained by heterogeneity in viscosity. (a) GEM particle mean squared displacement (MSD) vs time, calculated either by ensemble-averaging over all particle tracks (EA MSD, red line), or by first time-averaging over each track and then ensemble-averaging over all particles (TEA MSD, black line). 95% confidence intervals (CI) of the EA MSD were calculated by bootstrapping and are plotted as a red shaded region around the EA MSD. Note the logarithmic scale along the x- and y-axes. (b) The TEA and EA MSD calculated for a representative Doppelgänger simulation for Model #1: Uniform viscosity (see Fig. 3b). (c) The TEA and EA MSD calculated for a representative Doppelgänger simulation for Model #4: Spatial + cellular heterogeneity in viscosity (see Fig. 3b), using a 100 nm spatial domain size. (d-f) The percent difference between the EA and the TEA MSD ((EA-TEA)*100/EA) displayed in (a-c), respectively, plotted as a function of time interval. The best fit of the data to an exponential decay plus a constant: y = A*e(-Bt)+C is plotted as a thick dashed black line. (g) The percent difference between the EA and TEA MSD for Model #4: Spatial + cellular heterogeneity in viscosity, where each subplot represents a different choice for the domain size of the spatial heterogeneity. X- and y-axes for all subplots are identical. Each light orange line represents an individual simulation, equivalent to the entire experimental dataset. 50 replicate simulations are superimposed onto the plot. Each curve was individually fit to an exponential decay plus a constant: y = A*e(-Bt)+C, and the best fit parameters were averaged across all 50 simulations to produce the best fit line (thick dashed orange line). The best fit to the experimental data shown in (d) is overlaid as a thick dashed purple line. Of the domain sizes sampled, simulations using the 100 nm domain gives the closest agreement to the experimental data, with the experimental data best fit line lying well within the range of outcomes among replicate simulations. On average, the 100 nm simulation best fit line lies slightly below the experimental best fit line, and the 300 nm simulation best fit line lies slightly above the experimental best fit line. Thus we estimate the domain size of the cytoplasm is on the order of ~100-300 nm.

Another possible origin of non-ergodicity in particle diffusion is transient particle immobility, which can be mathematically described by a continuous time random walk (CTRW) (27). However, our longest-tracked diffusing particles (Fig. 1g) showed no evidence of transient immobilizations (Supp. Fig. 4a-b). Further, the time-averaged MSDs of individual tracks did not have a power law exponent, *α*, equal to 1 even for the longest-tracked particles (Supp. Fig. 4c), in contrast to what would be expected for a CTRW. Overall, our data were more consistent with heterogeneity in cytoplasm viscosity (Fig. 4) than a two-state system of mobile and immobile particles.

In the Doppelgänger simulations with heterogeneous viscosity, non-ergodicity arises from the fact that particles within distinct spatial domains exhibit different diffusion rates. Therefore, non-ergodicity in diffusion should depend on the domain size of the spatial heterogeneity in viscosity. Indeed, we found that non-ergodicity in our simulations is strongly dependent on the domain size (Fig. 4g). Interestingly, while the experimentally-measured *variability* in GEM diffusion can be reproduced using a wide range of different domain sizes (Table 4. Supp. Fig. 3), the experimentally-measured *non-ergodicity* of GEM particle diffusion could only be reproduced quantitatively with a subset of characteristic length scales (Fig. 4g). By comparing the simulations to the experimental data, we estimate the size of spatial domains of cytoplasm viscosity for the 40 nm GEM particles to be on the order of ~100-300 nm.

### Heterogeneity in diffusion does not arise from density fluctuations related to the cell cycle or cell tip growth

We next tested what factors might be responsible for such a large heterogeneity in cytoplasmic viscosity. Within an asynchronous population, fission yeast cells exhibit an approximately two-fold range in cell size, which corresponds to the cell cycle stage (63). A recent study used quantitative phase imaging (QPI) to show that the overall intracellular dry-mass density of fission yeast cells fluctuates over the cell cycle, with density decreasing during interphase and increasing during mitosis and cytokinesis (49). To test whether GEM diffusion also varies over the cell cycle, we examined the relationship of GEM diffusion with cell length as a proxy for cell cycle stage (Supp. Fig. 5a-b). We detected no significant correlation of diffusivity with cell length, making it unlikely that the cell-to-cell variability in GEM diffusion is cell cycle dependent.

We next tested whether spatial variations of density could explain the variability of GEM diffusion. QPI experiments demonstrated a subtle gradient of intracellular density in a subset of fission yeast cells, in which growing cell tips generally appear to be less dense than the rest of the cell (49). Regional cytoplasmic differences have also been shown in *Ashbya gossypii*, in which GEMs have decreased diffusivity in the perinuclear region (22). To test for spatial variations in fission yeast, we mapped the GEM tracks relative to their positions in the cell (Supp. Fig. 5c-d). This analysis yielded no obvious regional differences in diffusivity within the fission yeast cell; specifically, we noted no strong differences in diffusion at growing cell tips or at the perinuclear regions. Therefore, it is unlikely that systematic regional differences in intracellular density are responsible for the variance in diffusivity.

### Variance in diffusion is impacted by osmotic shock but not by cytoskeletal or temperature perturbations

We then probed what factors could affect the variance by submitting the cells to different perturbations. For each perturbation, we measured the distribution of track-wise and cell-wise fits of GEM diffusivities, and performed the Wilcoxon rank sum non-parametric test for equality of medians (64) and Levene’s test for equality of variances (60) to establish whether changes to the median and variance were statistically significant (Methods).

One cytoplasmic constituent implicated in the rheological properties of the cytoplasm is the cytoskeleton. A rigid and interconnected cytoskeleton network can act as a barrier (65), or elastically resist particle motion – properties which can be described by poroelastic models (40, 66). In addition, the cytoskeleton is responsible for transporting and positioning organelles and “actively mixes” the cytoplasm (67). The cytoskeleton may also create structured intracellular regions with distinct biophysical properties (68). We used a combination of latrunculin A (LatA) and methyl benzimidazol-2-yl-carbamate (MBC) to depolymerize actin and microtubules in interphase fission yeast cells (Fig. 5a). This treatment however had only subtle effects on GEM diffusivity; we detected a small, statistically insignificant increase in the median diffusivity (Fig. 5d, Wilcoxon t-test, Track-wise fits: 6% increase, p-value = 0.17; Cell-wise fits: 18% increase, p-value = 0.3), and a small, statistically insignificant increase in the variance (Fig. 5g,j, Levene test, Track-wise fits: 71% increase, p-value = 0.08; Cell-wise fits: 27% increase, p-value = 0.18)). We therefore concluded that the cytoskeleton is not the main determinant of cytoplasmic viscosity or variance at the 40-nm size scale in fission yeast.

**Figure 5:**
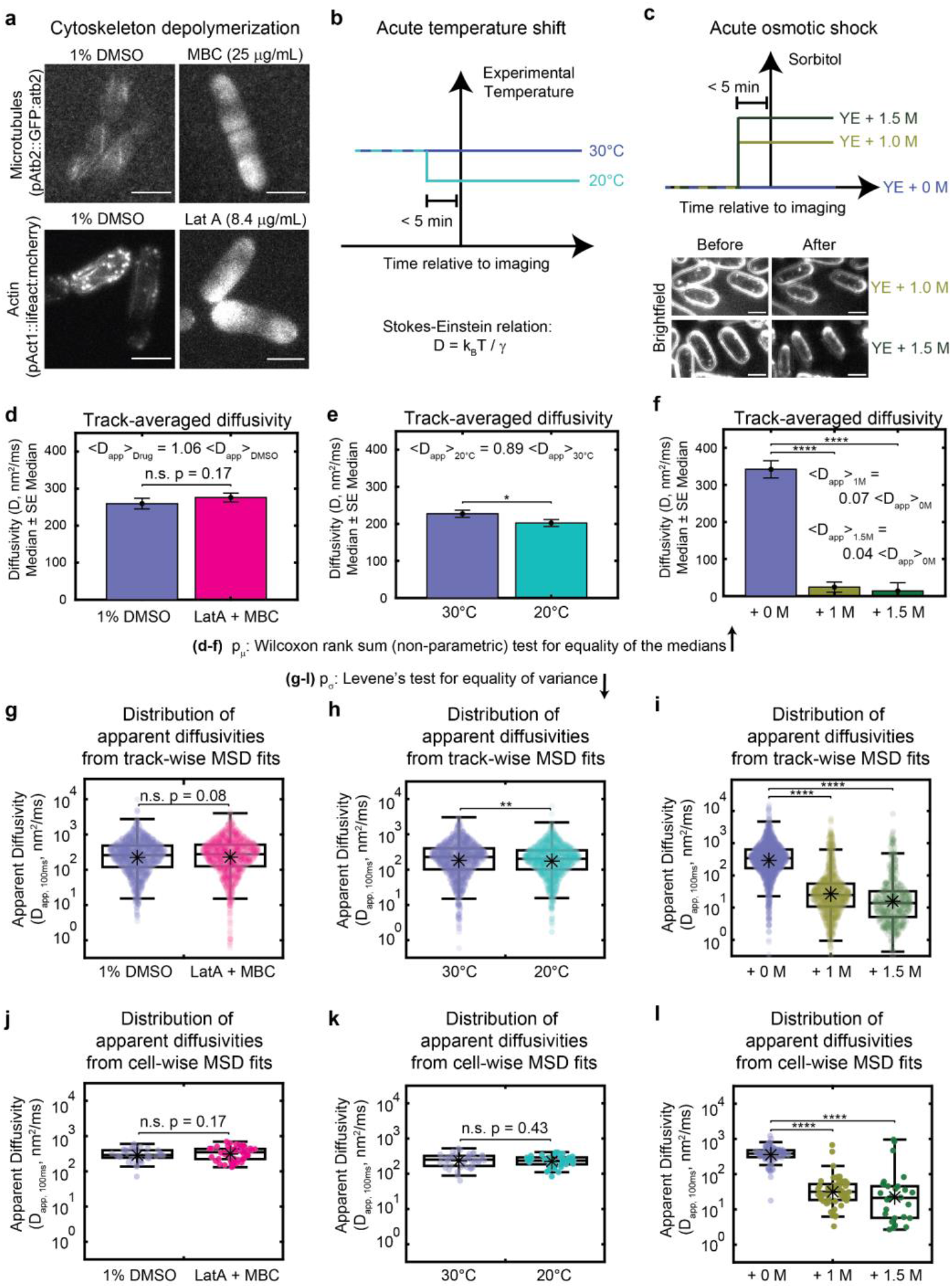
Heterogeneity in cytoplasmic diffusion has varied responsiveness to experimental perturbations. (**a**) Fluorescence images of fluorescent tubulin (top) and actin (bottom) in the context of the DMSO control (left) and addition of cytoskeleton depolymerizing drugs (right). Scale bar is 5 μm. (**b**) Schematic of experiments varying the experimental temperature (top) and prediction of the relationships between the diffusivity, *D*, and the experimental temperature, *T*, as well as the Boltzmann constant, *k_B_*, and the viscous drag coefficient, *γ* (bottom). (**c**) Schematic of experiments varying osmotic shock with sorbitol (top) and example brightfield images of osmotically-shocked cells showing a reduction in cell volume (bottom). Scale bar is 5 μm. (**d-f**) The median diffusivity is plotted for each experimental condition. Significance stars represent the result of the Wilcoxon rank sum test for equality of the medians. (**g-l**) Distributions of apparent diffusivities calculated from fits of the track-wise (**g-i**) or cell-wise (**j-l**) MSD curves displayed for each condition. Note the logarithmic scale along the y-axis. Boxplots are drawn as in Figure 2. Significance stars represent the result of Levene’s test for equality of variance. (**d-l**) * p<0.05. **p<0.01, *** p<0.001, **** p<0.0001.

Another main determinant of diffusivity *D* is temperature *T*. In addition to purely physical effects of temperature of diffusion as defined by the Stokes-Einstein equation where *D ∝ T*, temperature can also have also a multitude of biological effects. For instance, temperature shifts may alter active mixing of the cytoplasm (69), and trigger viscosity adaptation mechanisms via production of viscogens (70). There are also reports of regional differences in effective temperature within single cells (71–73).

To assay the effects of temperature on GEMs diffusivity, we grew fission yeast cells overnight at 30 °C, and then imaged them ~ 5 min after shifting cells down to 20 °C (Fig. 5b). This 10 °C decrease in temperature corresponds to ≈ 3% decrease in the absolute temperature (in Kelvin), and thus the Stokes-Einstein equation predicted a similar decrease in diffusivity. We observed a slightly larger than predicted drop in the median track-wise diffusivity of GEMs (Fig. 5e, Wilcoxon test, Track-wise fits: 11% decrease, p-value = 0.03; Cell-wise fits: 6% decrease, p-value = 0.54). The track-wise variance exhibited a statistically significant decrease, but the cell-wise variance did not change significantly (Levene test, Track-wise fits: 49% decrease, p-value = 0.006; Cell-wise fits: 28% decrease, p-value = 0.43). Overall, increasing the temperature had no effect on cell-to-cell variation but slightly increased intracellular heterogeneity.

Finally, we tested the effects of osmotic shocks. Osmotic shocks acutely alter the concentration of molecules in the cytoplasm by removal or addition of water (34, 37, 44). We performed hyperosmotic shocks with 1 M and 1.5 M sorbitol (Fig. 5c), which has been previously reported to roughly double the concentration of the cytoplasm (37, 44). Consistent with previous reports (37, 44), these hyper-osmotic shocks induced a striking decrease in the median track-averaged diffusivity of GEM particles compared to control experiments (Wilcoxon t-test, Track-wise fits: 93% and 96% decreases, p-values = 5*10^-273^, 3*10^-153^ for 1 M and 1.5 M shocks, respectively; Cell-wise fits: 92% and 94% decreases, p-values = 3*10^-21^, 4*10^-10^ for 1 M and 1.5 M, respectively). Interestingly, it also induced a sizable increase in both the track-wise and cell-wise variance in measured diffusivity (Fig. 5i,l, Levene test, Track-wise fits: 275% and 420% increases, p-values = 1*10^-9^, 2*10^-10^ for 1 M and 1.5 M shocks, respectively; Cell-wise fits: 530% and 16,083% increases, p-values = 4*10^-4^, 3*10^-9^ for 1 M and 1.5 M, respectively). Thus we found that increasing the concentration of the cytoplasm slowed diffusion but also drastically increased both intracellular and intercellular cytoplasmic heterogeneity. These results suggest that hyperosmotic shocks may make the cytoplasm even more heterogeneous.

## Discussion

Here we used a combined experimental and theoretical analysis to reveal a high degree of cytoplasmic heterogeneity experienced by objects on the scale of large protein complexes. In particular, our results indicated the effective cytoplasmic viscosity in fission yeast varies more than 10-fold among cells, and 100-fold within cells. Although the source of this heterogeneity is not yet understood, our analyses showed that viscosity variation is independent of the cytoskeleton, cell cycle stage, and temperature – but increases under hyperosmotic shock.

### Generalizability of cytoplasmic heterogeneity

It is highly likely that the large diffusive heterogeneity we observed in fission yeast is generalizable to most, if not all, cell types. In fact, because fission yeast exhibit strikingly regular cell shape and growth properties, they may be expected to have much less cytoplasmic variability than many other systems. Although most previous work has not explicitly focused on variability, studies of GEM particle diffusion in the cytoplasm or nucleoplasm of budding yeast, the filamentous fungus *Ashbya gossypii, Xenopus* egg extract, and several mammalian cell types (24, 26, 36, 41), as well as other studies of diffusion in *E. coli* (30), show that comparable variability in diffusion exists in these diverse contexts. In particular, McLaughlin et al. reported sizeable variation in both inter- and intra-cellular heterogeneity of GEM diffusivity in *Ashbya* (22). Beyond measurements of diffusion, a study directly probing viscosity also revealed substantial variability (74). Thus, large variability in cytoplasmic properties may be a fundamental, conserved property of cells. Hints from the literature suggest that heterogeneous cytoplasmic diffusion is also not limited to large protein complexes. Both larger objects such as lipid droplets (28, 54) and smaller particles such as individual fluorescent proteins (17–21, 23, 29, 75) and quantum dots (25, 26, 31) seem to exhibit substantial amounts of diffusive heterogeneity, as well as ergodicity-breaking (28–30).

### Sources of cytoplasmic heterogeneity

What might be the origin of this variability in cytoplasmic properties? Heterogeneity may originate from multiple non-exclusive sources. At the submicron and micron-scale, obstruction by organelles (76, 77) and other cytoplasmic structures such as condensates, as well as localized active mixing, could contribute to cytoplasmic variability (78). Local differences in the composition of specific macromolecules, for instance in the vicinity of organelles, may also contribute to cytoplasmic heterogeneity (79, 80). In addition, GEMs and other intracellular constituents may become transiently trapped between or inside membrane-bound and membrane-less compartments, thereby lowering the particle’s apparent diffusion rate. These various scenarios are consistent with our rough estimate of spatial domain size on the order of hundreds of nanometers for the 40 nm GEM particles.

At the nanometer-scale there are some enticing sources of heterogeneity that remain unexplored, notably those intrinsic to the macromolecular milieu: crowder density, size, charge, and hydrophobicity. Indeed, the fact that diffusion varies strongly with probe size and molecular species (25, 81–86), suggests that the local molecular structure of the cytoplasm plays a large role in the diffusion of macromolecules. Similarly, all-atom molecular dynamics simulations of the cytoplasm show that thermal fluctuations in the local cytoplasmic composition can lead to significant variability in diffusion rates (87). Therefore, the molecular and cellular features contributing to viscosity may themselves be highly dynamic and transient. Future studies of diffusive heterogeneity across different species, cell types, and physiological states will be invaluable for dissecting the biophysical determinants of cytoplasmic variation.

### Consequences of cytoplasmic heterogeneity

The heterogeneity of the cytoplasm may act as a highly significant source of biological noise for any diffusion-limited process. For example, spatial heterogeneity in diffusivity could lead to differences in diffusion-limited reaction rates across the cell. In particular, if the regions of high viscosity (low diffusivity) are long lived, they could act as “traps”, locally increasing the concentration of larger protein complexes or organelles, potentially influencing the speed and localization of certain reactions. The effects of stochasticity should be particularly strong for complexes which exist at low copy number or whose biological function depends on rare binding events. At the cell population level, having a wide range of diffusivities might be advantageous, allowing different cells to react to changes in the environment at different rates, permitting strategies such as bet-hedging to take place.

In fact, it is hard to imagine a biological process that would not be affected by such a large variation in the effective viscosity. For example, many reactions driving gene expression, biosynthesis and metabolism are considered to be diffusion limited. For example, cytoplasmic viscosity has been demonstrated to have strong effects on microtubule dynamics in vivo (37). Interestingly, the dynamics of individual microtubules were much less variable than those of the GEMs, suggesting that cellular systems may employ compensatory mechanisms that buffer the effects of heterogeneity in viscosity. Cellular control of viscosity and other aspects of the cytoplasm such as intracellular density represents a potential global mode of regulation.

### Generalization of the Doppelgänger simulation approach

Our analyses of biological noise were made possible by using our Doppelgänger simulation approach. This approach explicitly reproduces the experimental measurement statistics in silico, which allowed us to definitively distinguish between statistical noise and biological heterogeneity. This simulation approach may be generalizable to many other systems (Supp. Fig. 6), and could be useful for instance in the analysis of noise suppression. Overall, we believe this powerful approach combining experiment and theory will provide needed clarity for studies of stochastic processes in biology, such as cytoskeletal dynamics, signaling, and gene expression.

## Methods

### Yeast strains and culture conditions

Standard methods for growing and genetically manipulating *Schizosaccharomyces pombe* were used (89). The constructions of the GEMs expressing strains were described previously (37, 44). In brief, the encapsulin-mSapphire chimera was expressed under the control of the inducible *nmt1* promoter (90) on a multcopy pREP41X plasmid containing a leucine selection cassette. Cells were grown overnight in Edinburgh minimal medium (EMM) containing adenine, histidine and, uracil at 0.25 g per liter (here called EMM LEU-) and 0.1 μg/mL thiamine with shaking at 30 °C to exponential phase (OD600 between 0.2 - 0.8). See Table 1 for reagents and strain list. Expression of the *Pyrococcus furiosus* encapsulin-mSapphire construct produces particles of 40 nm in diameter, with the encapsulin proteins facing the inside of the particle and the fluorescent proteins facing the cytoplasm (36, 41).

### Microscopy

*S. pombe* cells were imaged in commercial microchannels (Ibidi μ-slide VI 0.4 slides; Ibidi 80606, Ibiditreat #1.5). Channels were pre-treated with 50 μl of 100 μg/ml lectin solution for 5 min. The lectin solution was removed by pipetting and 50 μl of cell culture were introduced then incubated for 5 to 10 minutes to allow adhesion to the lectin then cells were washed with EMM LEU-. For the 20°C condition the Ibidi slide and the buffer were equilibrated at 20°C before cells were added. For the 30°C condition, slides and buffers were equilibrated at 30°C before cells were added to it. For hyper-osmotic shocks, the medium was manually removed from the channel via pipetting and quickly replaced with pre-warmed (30°C) hyper-osmotic media. Cells were imaged immediately and for no longer than 5 minutes after the medium was exchanged to minimize adaptation. For cytoskeleton depolymerization cells were introduced in the Ibidi slide as described previously then the buffer was exchanged for prewarmed (30°C) EMM LEU-containing Latrunculin A (8.4 μg/mL or 20 μM) and methyl benzimidazol-2-yl-carbamate (MBC) (25 μg/mL or 131 μM). Cells were incubated at 30°C with the drug cocktail for 5 minutes prior to imaging. We confirmed that this treatment caused depolymerization of the microtubule and actin cytoskeletons in < 5 min by imaging cells expressing Lifeact-mCherry or GFP-Atb2.

For imaging GEMs, yeast cells were imaged with a Nikon TI-2 equipped with a Diskovery Multi-modal imaging system from Andor and a SCMOS camera (Andor, Ixon Ultra 888) using a 60x TIRF objective (Nikon, MRD01691). Cells were imaged sequentially, first a brightfield (BF) image then 1,000 fluorescence images at 100 Hz (for ~ 10 s) with a 488 nm excitation laser and a GFP emission filter 525 +/- 25 nm. Variable angle epifluorescence microscopy (VAEM) (51) was used to reduce background fluorescence and allow for the high imaging frequency required. Cells were selected for sparse numbers of labeled motile nanoparticles (< 10 GEMs per cell) to ensure proper particle tracking. Note that each GEM can be imaged multiple times during the acquisition giving more tracks per cell than the number of visible nanoparticles.

### Particle tracking

Cells were individualized from the field of view by cropping the images. Images of individual cells were rotated so that cell length (long axis) was horizontal. From the brightfield image cell length was measured by tracing a straight line joining each pole and passing through the center of the cell. Cell contours were drawn manually from the brightfield image and used to determine cell centroid. Cell length and centroid were used to plot GEMs tracks in linear and normalized space (Figure 1). GEMs nanoparticles in each cell were tracked using the MOSAIC plugin (Fiji ImageJ)(91, 92) with the following parameters for the 2D Brownian dynamics tracking in MOSAIC: radius = 3, cutoff = 0, per/abs = 0.2-0.3, link = 1, and displacement = 6. Tracks shorter than 10 timepoints were removed from further analysis.

### Diffusivity Analysis

Mean Square Displacement (MSD) Analysis: Unless explicitly stated otherwise, all analysis was performed using the time-ensemble averaged (TEA)*MSD = <(x(t+τ)-x(t))^2^>_t,n_*. The time-averaged MSD was first calculated individually for each track, and then a second averaging was performed to find the (ensemble averaged) MSD across all tracks. For the time-averaging of each track, MSDs were calculated using non-overlapping windows and plotted versus time interval, τ. For example, a track with 7 time steps and time interval τ = 3 time steps would have an MSD_x_ = ((x(t=4)-x(t=1))^2 + (x(t=7)-x(t=4))^2) / 2). For the subsequent ensemble-averaging step, the MSDs for each time interval were averaged across all tracks.

Fitting the MSD: A linear fit of *ln(MSD*) vs *ln(τ/τ_0_*) for the first 7 time intervals (~70 ms) was used to determine the values of the anomalous exponent and the apparent diffusivity (see our rationale for choice of *τ_0_* below in this paragraph). As the length of the trajectories is an exponentially decaying distribution (Fig. 1d - histograms), the statistical error grows with time (Fig. 1e - MSD) -- hence, we fit the only first part of the MSD function. The fitting resulted in two fit parameters corresponding to the equation *MSD = A(τ/τ_0_)^α^*, where *A* has units of nm^2^ (representing the MSD when *τ = τ_0_*) and *α* is unitless. We can convert these values to an apparent diffusivity by assuming *MSD_τ = τ0_* = *A* = *2nD_app,τ0_τ0*, where n is the number of spatial dimensions (in this case, n=2). Solving for the apparent diffusivity *D_app_* in nm^2^ms^-1^, we find the following conversion: *D_app,τ0_* = *A/(2nτ_0_*) (representing the apparent diffusivity specifically at *τ_0_*). We choose *τ_0_* = 100ms, to represent the intermediate regime measured in our dataset. For track-wise fits, the time-averaged MSD was calculated and fit separately for each trajectory. For cell-wise and condition-wise MSD calculations, the time-averaged MSDs for each track were then ensemble-averaged over all tracks in each cell or condition, respectively, and subsequently fit. The 95% confidence intervals (CI) for *α* and *D_app_* were calculated using bootstrapping of the TEA MSD by sampling the individual TA MSDs; the bootci() function in MATLAB was called using the basic percentile method and a sample size equal to the number of tracks in the dataset.

### Doppelganger Simulations

Simulations of particle diffusion were implemented using fixed time step Brownian dynamics, according to the Stokes-Einstein relation for diffusion of a spherical particle in a viscous medium (*D = k_B_T/γ*, where *D* is the diffusion coefficient such that the mean-squared displacement *MSD = 2nDt* is linear with time *t* and the number of dimensions *n*, *kB* is the Boltzmann constant, *T* is the temperature, and *γ = 6πηR*, where *η* is the viscosity of the cytoplasm and *R* is the radius of the particle). See Tables 2–3 for a list of the parameters used. All code was written in custom MATLAB scripts. Cells were implemented as 2D rectangular boxes with reflecting boundary conditions at the edges of each box. All simulated cells had a short-axis width of 3 μm, and a long-axis width equal to that of its experimentally-measured doppelgänger. (Note that the short-axis width was chosen to be 3 μm, rather than the known 4 μm diameter of fission yeast cells, to best represent the imaging conditions in the experimental data. VAEM imaging only captures the lower portion of the cell near the coverslip, where the cross-section is smaller than at the equatorial plane.)

Each simulated cell had the same number of particles as its experimental doppelgänger. Each particle was initialized randomly within the rectangular cell wall boundary. After initialization, particle positions were updated using fixed time step Brownian dynamics, where the fixed time step, *Δt*, was equal to the acquisition frame rate of the experimental measurements. In each time step, a random number generator (randn, seeded randomly at the beginning of each set of simulations with rng(‘shuffle’)) selected each particle’s step size and direction from a normal distribution with a mean of zero and a standard deviation of ξ = sqrt(2*k_B_T\**Λ/γ*). If a particle left the cell boundary during a timestep, the particle’s position was reflected across the cell boundary (or boundaries) that the particle crossed, in order to keep the particle inside the cell (i.e., reflective boundary conditions). For ease of implementation, all particle tracks were simulated for the longest length of time any particle in the experimental dataset was tracked; then after simulations were complete, each simulated particle’s data were pruned to match their experimental doppelgänger -- all other timepoints that were not tracked for the experimental doppelgänger were deleted from the simulated dataset.

For simulations with cell-cell variations in viscosity (Models #3 and #4), viscosity values for each cell were chosen from a random log-normal distribution (*η = μe^(σ/μ)*randn()^*), with a mean viscosity equal to 40 times that of water, and a standard deviation of 45% of the mean. For simulations with spatially-varying viscosity (Models #2 and #4), each rectangular cell was broken up into spatial domains of equally-sized squares with 1 μm side-length. As all cells were 3 μm in width but variable in length, simulated spatial domains within cells were arranged in a 3xm grid, where m is the number of domains along the long axis. If the cell length along the long dimension was not an integer multiple of 1 μm, then the remainder was placed in its own spatial domain of smaller size. Viscosity values in each domain were chosen from a random log-normal distribution (*η = μe^(σ/μ)*randn()^*), with a mean equal to the mean viscosity of that cell, and a standard deviation equal to 85% of the mean.

For simulations varying the domain size of spatial heterogeneity, the mean and variance in viscosity was fit to the experimental data in order to replicate the mean and variance of the experimentally-measured GEM particle diffusivity (Supp. Fig. 4, Table 4). The ergodicity was then compared between different spatial domain sizes under these conditions.

### Statistical analysis

Velocity autocorrelation analysis: Velocity autocorrelations were defined as *VAC(τ) = <(v(t+τ)v(t))> t* and were performed using non-overlapping intervals.

ANOVA: A nested, n-way analysis of variance was performed using MATLAB’s anovan() function. Track identity was nested under cell, session, and day identities, cell identity was nested under session and day identities, and the session identity was nested under the day identity. ANOVA was performed separately on the power law exponents and the natural logarithm of the diffusivities. ANOVA was performed identically on the experimental and simulated datasets. Because the Doppelgänger simulation approach computationally reproduces the exact experimental distribution of tracks, cells, sessions, and days, the exact magnitudes of the variance attributed to each category can be directly quantitatively compared (e.g. Fig. 3f).

Comparison of median diffusivity values between conditions: A Wilcoxson rank sum non-parametric test for equality of medians (64) was performed to determine whether differences in the medians between conditions were statistically significant. We chose a non-parametric test, and compared the medians instead of the means, so that our analysis would be less sensitive to the fact that the distributions were long-tailed and not perfectly Gaussian (even on a log scale). Statistical tests were performed on the logarithm (base 10) of the apparent diffusivities.

Comparison of variance in diffusivity values between conditions: Levene’s test for equality of variance (60) was performed to determine whether differences in the variances between conditions were statistically significant. While Levene’s test is not a non-parametric test, it is less sensitive to non-normality than many other parametric tests, and is MATLAB’s recommended test for equality of variance for non-normal distributions. Statistical tests were performed on the logarithm (base 10) of the apparent diffusivities.

Converting summary statistics from log space to linear space: Because the diffusivities were more normally distributed on a log scale than a linear scale, all of the summary statistics (medians, standard deviations, etc.) were calculated on the distribution in log space. For a distribution that is log-normally distributed, medians and standard deviations calculated in log space are not the same as those calculated in linear space and so are not interchangeable (i.e. 10^<a>^ =/= <10^a^>, and 10^sqrt(<(a-<a>)^2>)^ =/= sqrt(<(10^a^-<10^a^>)^2^>) and have different interpretations. The median of the log-scale diffusivity distribution (μlog = <log_10_(D_app,100ms_)> represents the median order of magnitude of diffusivities in the dataset.The medians reported in this work were first calculated from the distribution in log space, and then converted to linear space as μ_linear_ = 10^μ_log^, and also represents the median order of magnitude (but now presented in linear space). The standard deviation of the log-scale diffusivity distribution (σ = sqrt(<(log_10_(D_app,100ms_)-<log_10_(D_app,100ms_)>)^2>) represents the number of orders of magnitude spanned by the dataset). In linear space, the associated number which best captures the data’s span in order of magnitudes is the fold-range of the distribution measured at some specified number of standard deviations away from the mean. In our dataset, a 2.5σ threshold best matched the outlier exclusion algorithm used in our box-plotting software (1.5 times the interquartile range past the 25th and 75th percentiles, Fig. 2c, see caption). To determine the fold-range, the standard deviation was calculated for the diffusivity distribution in log space, then the ratio of the diffusivities at 10^(μ-log±2.5*σ_log)^ (i.e., the fold-range) was evaluated as 10^(μ+2.5*σ)^/10^(μ-2.5*σ)^ = 10^5*σ^. In perturbation conditions (Fig. 5), the reported percent change in the median and fold-range were determined using the converted linear space median and fold-range as described above in this paragraph.

### Ergodicity

The ensemble-average (EA)*MSD = <(x(τ)-x(0))^2^>* was computed as the squared displacement at each time interval relative to the particle’s origin position, and then averaged across all particles. The 95% confidence intervals (CI) for the EA MSD were calculated using bootstrapping; the bootci() function in MATLAB was called using the basic percentile method and a sample size equal to the number of tracks in the dataset. The time-ensemble-average (TEA) *MSD = <(x(t+τ)-x(t))^2^>* was first time-averaged across each individual track using non-overlapping time intervals, and then the time-averages were again averaged across all particles (exactly as in *Diffusivity Analysis*). The percent difference between the EA and TEA MSD was calculated as (EA-TEA)*100/EA MSD. The percent difference as a function of the time interval was then fit to an exponential decay plus a constant: y = A*e(-Bt)+C using the MATLAB fit() function. The fitting was weighted by the inverse of the standard error for each data point.

## Data and code availability

All raw imaging data are available upon request. All tracking data and code are freely available on Gitlab: <https://gitlab.com/theriot_lab/vast-heterogeneity-in-cytoplasmic-diffusion-rates-revealed-by-nanorheology-and-doppelgaenger-simulations.git>

## Author Contributions

Conceptualization, A.T.M., and R.M.G.;

Methodology, A.T.M, R.M.G.;

Software, R.M.G.;

Validation, R.M.G. and A.T.M.;

Formal Analysis, R.M.G., and A.T.M.;

Investigation, A.T.M., and R.M.G;

Resources, A.T.M., and R.M.G.;

Data curation, A.T.M. and R.M.G.;

Writing – Original Draft, A.T.M., R.M.G.,;

Writing – Review & Editing, A.T.M., R.M.G., J.A.T., and F.C.;

Visualization, R.M.G. and A.T.M.;

Supervision, A.T.M. and R.M.G.;

Project Administration, A.T.M. and R.M.G.;

Funding Acquisition, J.A.T. and F.C.;

R.M.G. and A.T.M. contributed equally and have the right to list their name first on their C.V.

## Acknowledgements

We thank Joël Lemière and Elena Koslover, for useful discussions and Chenlei Hu, for preliminary analysis and modeling. We are grateful to the MBL community and members of the MBL Physical Biology of the Cell Course for support. We thank the Chang lab and the Theriot labs for support and discussions.

## Funding

This work was supported by grants to F.C. (NIH R01GM115185 and R35GM141796), grants to J.A.T. (NIH R37-AI036929 and the Howard Hughes Medical Institute), a Gerald J. Lieberman Fellowship to R.M.G, and an NSF Graduate Research Fellowship to R.M.G.

This article is subject to HHMI’s Open Access to Publications policy. HHMI lab heads have previously granted a nonexclusive CC BY 4.0 license to the public and a sublicensable license to HHMI in their research articles. Pursuant to those licenses, the author-accepted manuscript of this article can be made freely available under a CC BY 4.0 license immediately upon publication.

## Supplementary Figures

**Supp. Fig. 1:**
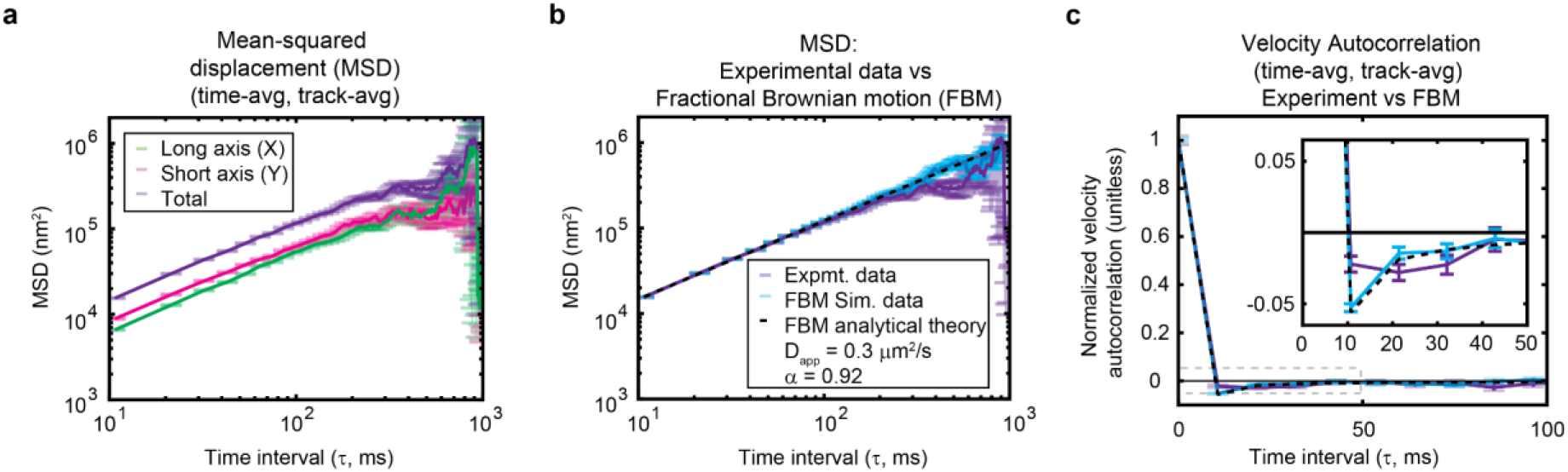
Experimental data is consistent with nearly unconstrained diffusion. (**a**) Mean-squared displacement (MSD) along the long and short axes of the cell, plotted alongside the total MSD. (**b**) The predicted MSD for Fractional Brownian motion (FBM), including both analytical theory and results from simulated data, using the experimentally-measured values of *D* and α. The experimental data is also plotted for comparison, showing good agreement with the theory. (**a-b**) Note the logarithmic scale along the x- and y-axes. (**c**) Same legend as in (b). The predicted velocity autocorrelation for FBM, showing the characteristic negative peak which then decays to zero. Experimental data shows a wide and very shallow negative basin, which does not match the shape or depth of the peak predicted by FBM.

**Supp. Fig. 2:**
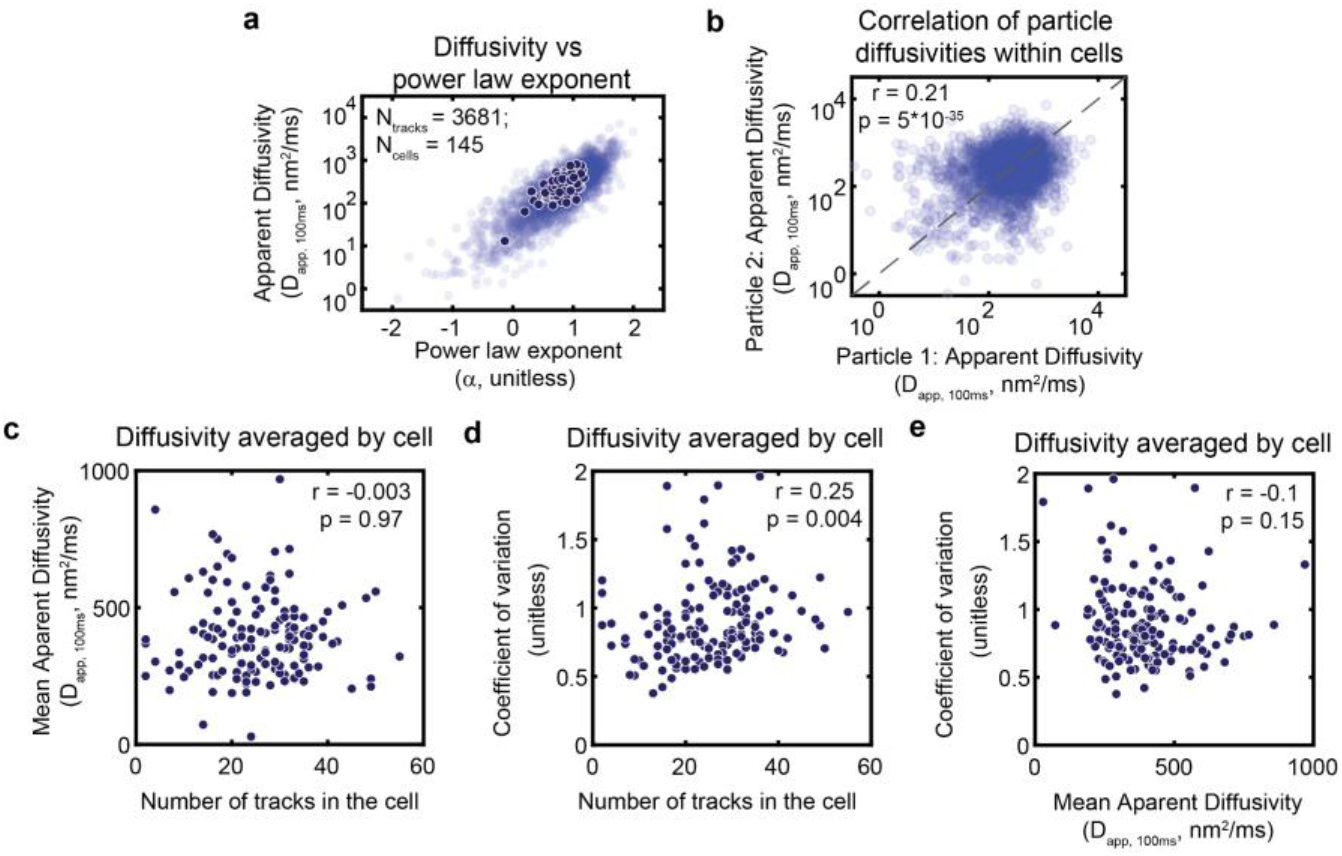
Additional evidence for intrinsic and extrinsic sources of noise. (**a**) The relationship between the apparent diffusivity and power law exponent. (**b**) The apparent diffusivity of each particle in the dataset plotted against a randomly chosen particle from the same cell. Each particle is represented exactly once in the plot. For cells with an odd number of particles, one particle would not be represented for that cell. (**c**) Mean diffusivity across tracks in each cell plotted vs the number of tracks in each cell. (**d**) Coefficient of variation across tracks in each cell plotted vs the number of tracks in each cell. (**e**) Coefficient of variation vs mean diffusivity calculated by averaging across all tracks for each cell. (**a-e**) Fits of track-wise MSD data are shown in light blue, with cell-wise fits overlaid in dark blue. (**a-b**) Note the logarithmic scale along the y-axis. (**b**) Note the logarithmic scale along the x-axis. (**b-d**) r- and p-values determined by a Spearman correlation algorithm.

**Supp. Fig. 3:**
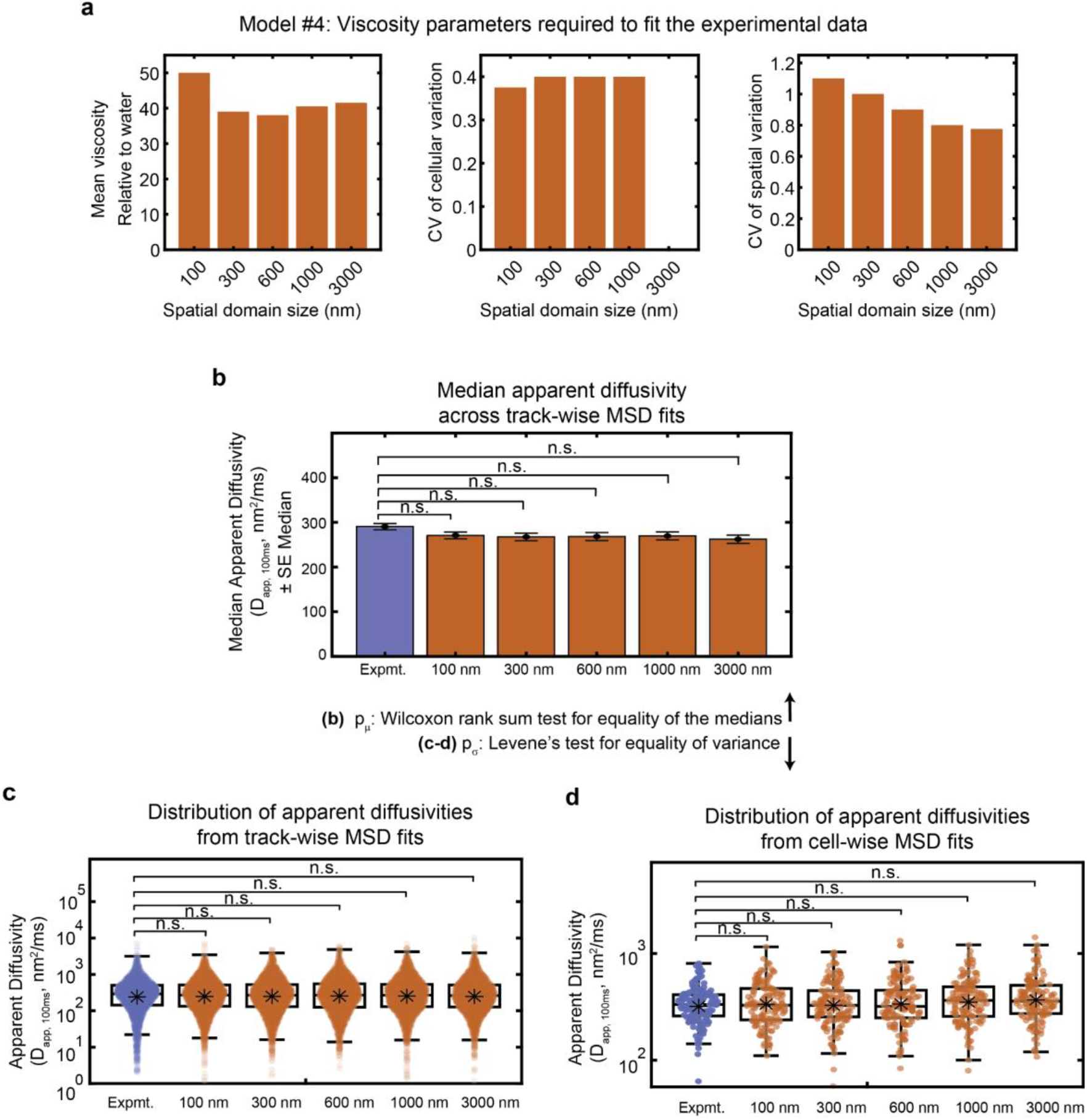
Best fit parameters for each spatial domain size preserve the experimentally-observed mean and variance in diffusivity. (**a**) Simulation input parameters for viscosity (Model #4: Spatial and cellular heterogeneity) that best recapitulate the experimentally-measured spread in diffusivity. *Left:* The mean viscosity relative to the viscosity of water (e.g., A mean of 40 would indicate the cytoplasm has 40X the viscosity of water). *Middle:* The coefficient of variation (CV, mean divided by the standard deviation) of the viscosity among different spatial domains within each cell. *Right:* The coefficient of variation (CV, mean divided by the standard deviation) of the cell-averaged viscosities among a population of cells. (**b**) Median apparent diffusivity (averaged across all tracks) plotted for the experimental dataset as well as each model. X-labels for the models represent the domain size for the spatial heterogeneity. Error bars represent the standard error of the median. Significance stars represent the result of the Wilcoxon rank sum test for equality of the medians. (**c-d**) Distributions of apparent diffusivities calculated from fits of the track-wise (**c**) or cell-wise (**d**) MSD curves displayed for the experimental data as well as each of the models. Note the logarithmic scale along the y-axis. Boxplots are drawn as in Figure 2. Significance stars represent the result of Levene’s test for equality of variance. (**a-c**) * p<0.05. **p<0.01, *** p<0.001, **** p<0.0001.

**Supp. Fig. 4:**
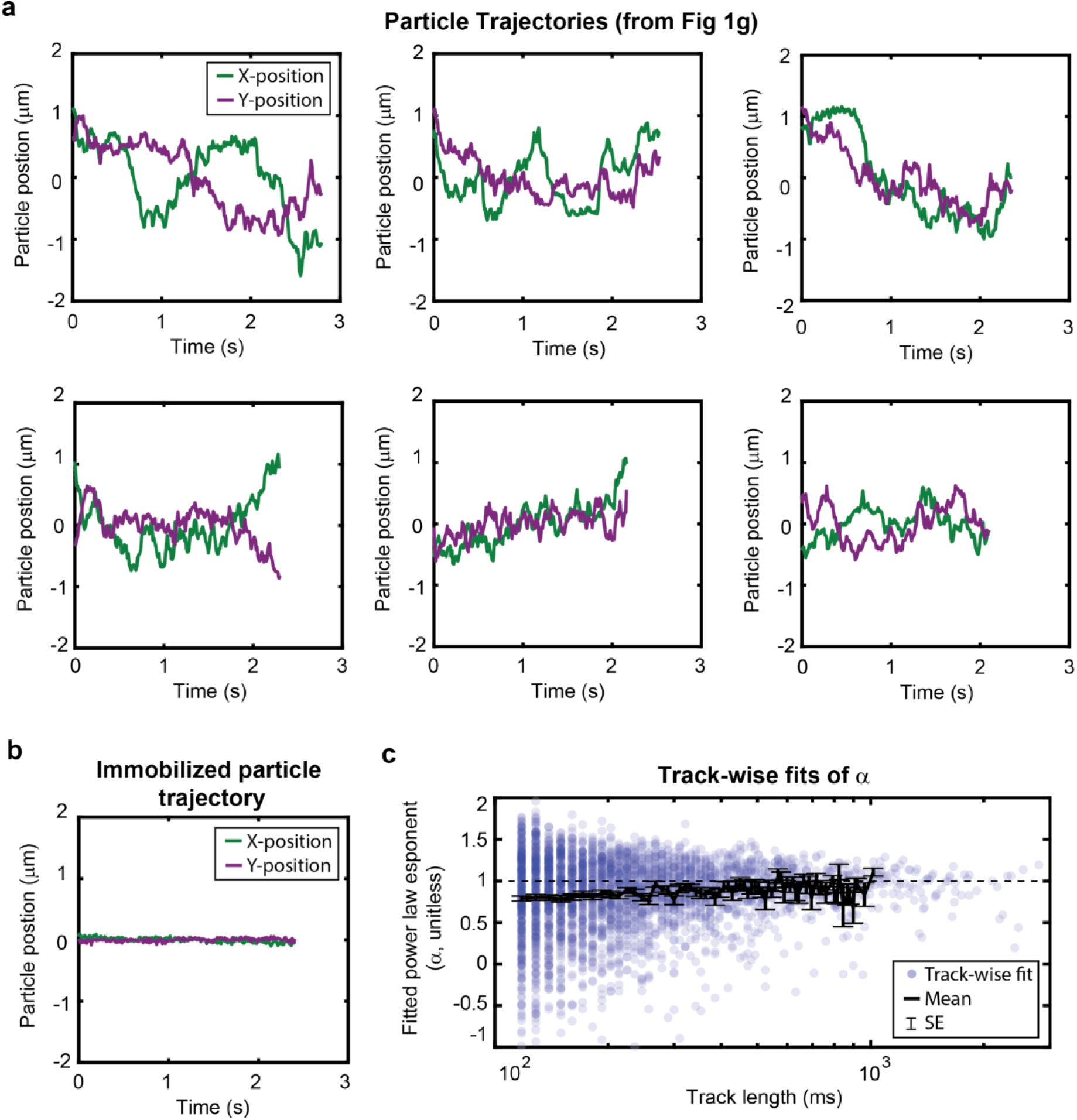
Weak non-ergodicity of GEM diffusion cannot be explained by a continuous time random walk model. (**a**) X- and y-trajectories of the tracks shown in Fig. 1g. (**b**) X- and y-trajectories of a completely immobilized particle observed within the experimental dataset. (**c**) The best fit of the power law exponent, α, for time-averaged MSD of each track, plotted as a function of the track length. Each dot represents the best fit for an individual track. The mean across all tracks of a given length is displayed as a thick black line, and the standard error of the mean (SE) is plotted as error bars.

**Supp. Fig. 5:**
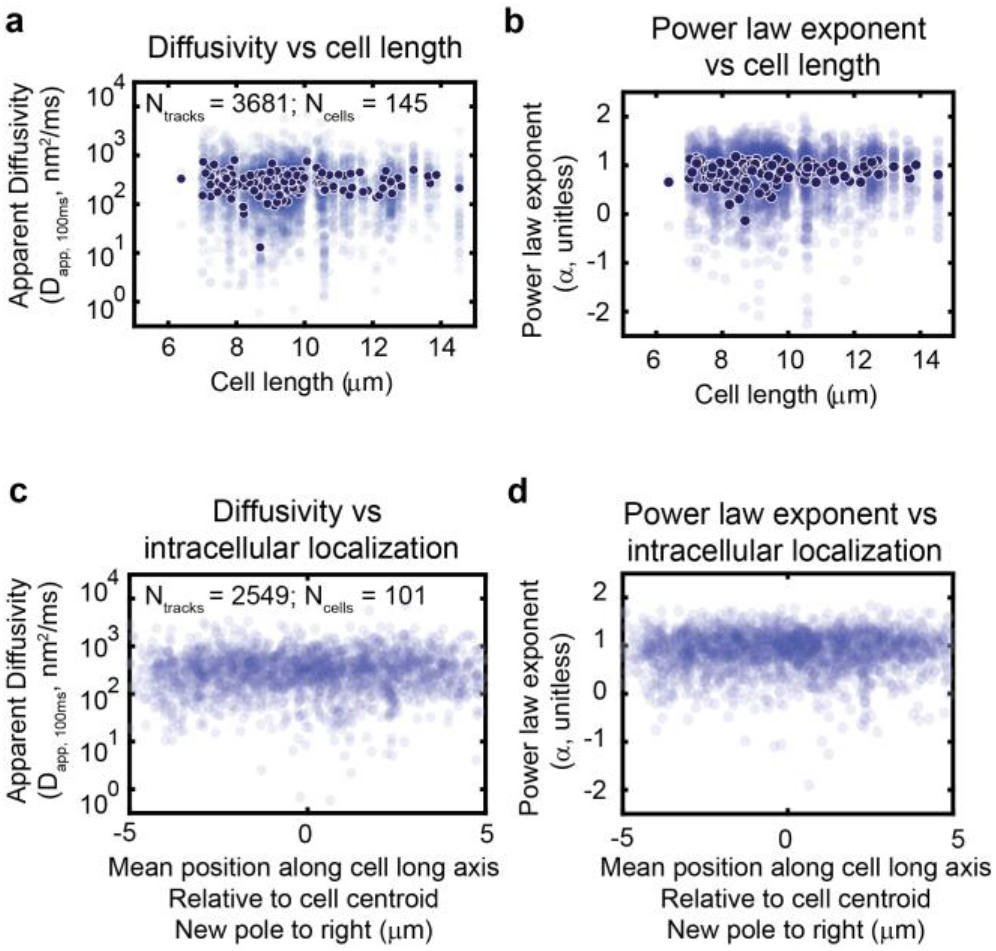
The large heterogeneity in diffusivity cannot be explained by the cell cycle or subcellular GEM particle localization. (**a-b**) Fitted values for diffusivity (a) and power law exponent (**b**) plotted as a function of cell length. (**c-d**) Track-wise fit values for diffusivity (**c**) and power law exponent (**d**) plotted against the mean (time-averaged) particle position along the long axis of the cell. There are fewer cells and tracks represented in (**c-e**) compared to (**a-b**) because the new pole could be distinguished from the old pole for only a subset of cells. (**a-d**) Fits of track-wise MSD data are shown in light blue, with cell-wise fits overlaid in dark blue. (**a, c**) Note the logarithmic scale along the y-axis.

**Supp. Fig. 6:**
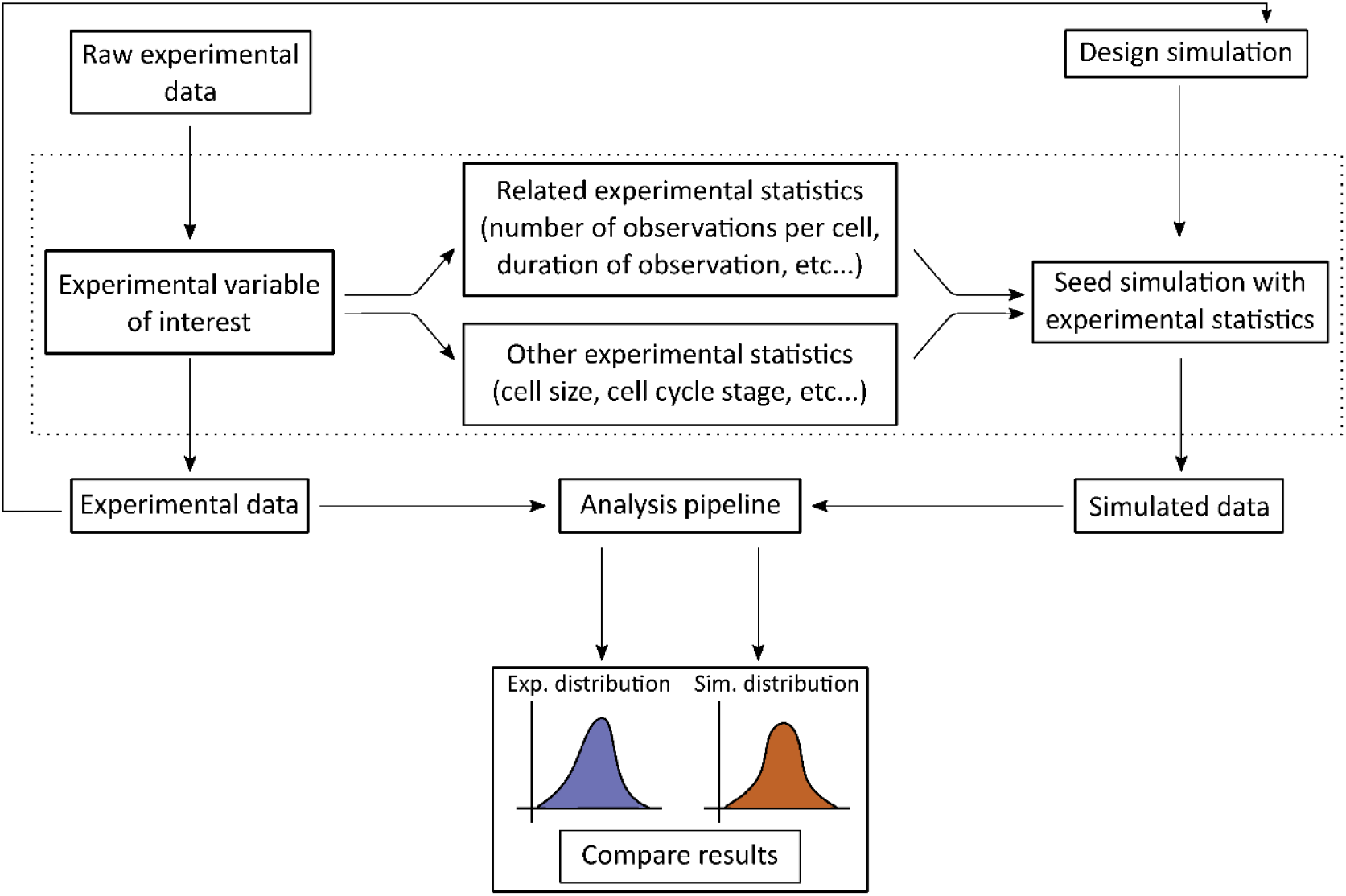
Schematic of the generalized Doppelgänger approach.

## References

1. Huang, W.Y.C., Q. Yan, W.-C. Lin, J.K. Chung, S.D. Hansen, S.M. Christensen, H.-L. Tu, J. Kuriyan, and J.T. Groves. 2016. Phosphotyrosine-mediated LAT assembly on membranes drives kinetic bifurcation in recruitment dynamics of the Ras activator SOS. Proc. Natl. Acad. Sci. U. S. A. 113:8218–8223.

2. Schmoller, K.M. 2017. The phenomenology of cell size control. Curr. Opin. Cell Biol.49:53–58.

3. Facchetti, G., B. Knapp, F. Chang, and M. Howard. 2019. Reassessment of the Basis of Cell Size Control Based on Analysis of Cell-to-Cell Variability. Biophys. J. 117:1728–1738.

4. Patterson, J.O., P. Rees, and P. Nurse. 2019. Noisy Cell-Size-Correlated Expression of *Cyclin B Drives Probabilistic Cell-Size Homeostasis in Fission Yeast*. Curr. Biol.29:1379–1386.e4.

5. Chang, A.Y., and W.F. Marshall. 2017. Organelles--understanding noise and heterogeneity in cell biology at an intermediate scale. J. Cell Sci. 130:819–826.

6. Mohapatra, L., B.L. Goode, P. Jelenkovic, R. Phillips, and J. Kondev. 2016. Design Principles of Length Control of Cytoskeletal Structures. Annu. Rev. Biophys. 45:85–116.

7. Gray, W.T., S.K. Govers, Y. Xiang, B.R. Parry, M. Campos, S. Kim, and C. Jacobs-Wagner. 2019. Nucleoid Size Scaling and Intracellular Organization of Translation across Bacteria. Cell. 177:1632–1648.e20.

8. Oates, A.C. 2011. What’s all the noise about developmental stochasticity? Development. 138:601–607.

9. Raj, A., and A. van Oudenaarden. 2008. Nature, nurture, or chance: stochastic gene expression and its consequences. Cell. 135:216–226.

10. Raser, J.M., and E.K. O’Shea. 2005. Noise in Gene Expression: Origins,Consequences, and Control. Science.

11. Battich, N., T. Stoeger, and L. Pelkmans. 2015. Control of Transcript Variability in Single Mammalian Cells. Cell. 163:1596–1610.

12. Suderman, R., J.A. Bachman, A. Smith, P.K. Sorger, and E.J. Deeds. 2017. Fundamental trade-offs between information flow in single cells and cellular populations. Proc. Natl. Acad. Sci. U. S. A. 114:5755–5760.

13. Levien, E., J. Min, J. Kondev, and A. Amir. 2021. Non-genetic variability in microbial populations: survival strategy or nuisance? Rep. Prog. Phys. 84.

14. Milo, R., and R. Phillips. 2015. Cell Biology by the Numbers. Garland Science.

15. Requião, R.D., L. Fernandes, H.J.A. de Souza, S. Rossetto, T. Domitrovic, and F.L. Palhano. 2017. Protein charge distribution in proteomes and its impact on translation. PLoS Comput. Biol. 13:e1005549.

16. White, S.H., and R.E. Jacobs. 1990. Statistical distribution of hydrophobic residues along the length of protein chains. Implications for protein folding and evolution. Biophys. J. 57:911–921.

17. Bakshi, S., B.P. Bratton, and J.C. Weisshaar. 2011. Subdiffraction-limit study of Kaede diffusion and spatial distribution in live Escherichia coli. Biophys. J. 101:2535–2544.

18. Baum, M., F. Erdel, M. Wachsmuth, and K. Rippe. 2014. Retrieving the intracellular topology from multi-scale protein mobility mapping in living cells. Nat. Commun. 5:4494.

19. Dross, N., C. Spriet, M. Zwerger, G. Müller, W. Waldeck, and J. Langowski. 2009. Mapping eGFP oligomer mobility in living cell nuclei. PLoS One. 4:e5041.

20. Manley, S., J.M. Gillette, G.H. Patterson, H. Shroff, H.F. Hess, E. Betzig, and J. Lippincott-Schwartz. 2008. High-density mapping of single-molecule trajectories with photoactivated localization microscopy. Nat. Methods. 5:155–157.

21. Scipioni, L., M. Di Bona, G. Vicidomini, A. Diaspro, and L. Lanzanò. 2018. Local raster image correlation spectroscopy generates high-resolution intracellular diffusion maps. Commun Biol. 1:10.

22. McLaughlin, G.A., E.M. Langdon, J.M. Crutchley, L.J. Holt, M.G. Forest, J.M. Newby, and A.S. Gladfelter. 2020. Spatial heterogeneity of the cytosol revealed by machine learning-based 3D particle tracking. Mol. Biol. Cell. 31:1498–1511.

23. Xiang, L., K. Chen, R. Yan, W. Li, and K. Xu. 2020. Single-molecule displacement mapping unveils nanoscale heterogeneities in intracellular diffusivity. Nat. Methods.17:524–530.

24. Huang, W.Y.C., X. Cheng, and J.E. Ferrell. 2021. Cytoplasmic organization promotes protein diffusion. bioRxiv. 2021.07.09.451827.

25. Etoc, F., E. Balloul, C. Vicario, D. Normanno, D. Liße, A. Sittner, J. Piehler, M. Dahan,and M. Coppey. 2018. Publisher Correction: Non-specific interactions govern cytosolic diffusion of nanosized objects in mammalian cells. Nat. Mater. 17:1048.

26. Sabri, A., X. Xu, D. Krapf, and M. Weiss. 2020. Elucidating the Origin of Heterogeneous *Anomalous Diffusion in the Cytoplasm of Mammalian Cells*. Phys. Rev. Lett.125:058101.

27. Lubelski, A., I.M. Sokolov, and J. Klafter. 2008. Nonergodicity mimics inhomogeneity in single particle tracking. Phys. Rev. Lett. 100:250602.

28. Jeon, J.-H., V. Tejedor, S. Burov, E. Barkai, C. Selhuber-Unkel, K. Berg-Sørensen, L. Oddershede, and R. Metzler. 2011. In vivo anomalous diffusion and weak ergodicity breaking of lipid granules. Phys. Rev. Lett. 106:048103.

29. Weigel, A.V., B. Simon, M.M. Tamkun, and D. Krapf. 2011. Ergodic and nonergodic processes coexist in the plasma membrane as observed by single-molecule tracking. Proc. Natl. Acad. Sci. U. S. A. 108:6438–6443.

30. Parry, B.R., I.V. Surovtsev, M.T. Cabeen, C.S. O’Hern, E.R. Dufresne, and C. Jacobs-Wagner. 2014. The bacterial cytoplasm has glass-like properties and is fluidized by metabolic activity. Cell. 156:183–194.

31. Janczura, J., M. Balcerek, K. Burnecki, A. Sabri, M. Weiss, and D. Krapf. 2021. Identifying heterogeneous diffusion states in the cytoplasm by a hidden Markov model. New J. Phys. 23:053018.

32. Cadart, C., L. Venkova, P. Recho, M.C. Lagomarsino, and M. Piel. 2019. The physics of cell-size regulation across timescales. Nat. Phys. 15:993–1004.

33. Neurohr, G.E., and A. Amon. 2020. Relevance and Regulation of Cell Density. Trends Cell Biol. 30:213–225.

34. Knapp, B.D., P. Odermatt, E.R. Rojas, W. Cheng, X. He, K.C. Huang, and F. Chang. 2019. Decoupling of Rates of Protein Synthesis from Cell Expansion Leads to Supergrowth. Cell Syst. 9:434–445.e6.

35. Tsai, H.-J., A.R. Nelliat, M.I. Choudhury, A. Kucharavy, W.D. Bradford, M.E. Cook, J. Kim, D.B. Mair, S.X. Sun, M.C. Schatz, and R. Li. 2019. Hypo-osmotic-like stres sunderlies general cellular defects of aneuploidy. Nature. 570:117–121.

36. Delarue, M., G.P. Brittingham, S. Pfeffer, I.V. Surovtsev, S. Pinglay, K.J. Kennedy, M. Schaffer, J.I. Gutierrez, D. Sang, G. Poterewicz, J.K. Chung, J.M. Plitzko, J.T. Groves, C. Jacobs-Wagner, B.D. Engel, and L.J. Holt. 2018. mTORC1 Controls Phase Separation and the Biophysical Properties of the Cytoplasm by Tuning Crowding. Cell.174:338–349.e20.

37. Molines, A.T., J. Lemière, M. Gazzola, I.E. Steinmark, C.H. Edrington, C.-T. Hsu, P. Real-Calderon, K. Suhling, G. Goshima, L.J. Holt, M. Thery, G.J. Brouhard, and F. Chang. 2022. Physical properties of the cytoplasm modulate the rates of microtubule polymerization and depolymerization. Dev. Cell. 57:466–479.e6.

38. Neurohr, G.E., R.L. Terry, J. Lengefeld, M. Bonney, G.P. Brittingham, F. Moretto, T.P. Miettinen, L.P. Vaites, L.M. Soares, J.A. Paulo, J.W. Harper, S. Buratowski, S. Manalis, F.J. van Werven, L.J. Holt, and A. Amon. 2019. Excessive Cell Growth Causes Cytoplasm Dilution And Contributes to Senescence. Cell. 176:1083–1097.e18.

39. Guo, M., A.F. Pegoraro, A. Mao, E.H. Zhou, P.R. Arany, Y. Han, D.T. Burnette, M.H. Jensen, K.E. Kasza, J.R. Moore, F.C. Mackintosh, J.J. Fredberg, D.J. Mooney, J. Lippincott-Schwartz, and D.A. Weitz. 2017. Cell volume change through water efflux impacts cell stiffness and stem cell fate. Proc. Natl. Acad. Sci. U. S. A. 114:E8618–E8627.

40. Charras, G.T., T.J. Mitchison, and L. Mahadevan. 2009. Animal cell hydraulics. J. Cell Sci. 122:3233–3241.

41. Szórádi, T., T. Shu, G.R. Kidiyoor, Y. Xie, N.L. Herzog, A. Bazley, M. Bonucci, S. Keegan, S. Saxena, F. Ettefa, G. Brittingham, J. Lemiere, D. Fenyö, F. Chang, M. Delarue, and L.J. Holt. 2021. nucGEMs probe the biophysical properties of the nucleoplasm. bioRxiv. 2021.11.18.469159.

42. Carlini, L., G.P. Brittingham, L.J. Holt, and T.M. Kapoor. 2020. Microtubules Enhance Mesoscale Effective Diffusivity in the Crowded Metaphase Cytoplasm. Dev. Cell.54:574–582.e4.

43. Molines, A.T., J. Lemière, M. Gazzola, I.E. Steinmark, C.H. Edrington, C.-T. Hsu, P. Real-Calderon, K. Suhling, G. Goshima, L.J. Holt, M. Thery, G.J. Brouhard, and F. Chang. 2022. Physical properties of the cytoplasm modulate the rates of microtubule polymerization and depolymerization. Dev. Cell. 57:466–479.e6.

44. Lemière, J., P. Real-Calderon, L.J. Holt, T.G. Fai, and F. Chang. 2022. Control of nuclear size by osmotic forces in Schizosaccharomyces pombe. Elife. 11.

45. Alric, B., C. Formosa-Dague, E. Dague, L.J. Holt, and M. Delarue. 2021. Macromolecular crowding limits growth under pressure. bioRxiv. 2021.06.04.446859.

46. Saunders, T.E., K.Z. Pan, A. Angel, Y. Guan, J.V. Shah, M. Howard, and F. Chang. 2012. Noise reduction in the intracellular pom1p gradient by a dynamic clustering mechanism. Dev. Cell. 22:558–572.

47. Abenza, J.F., E. Couturier, J. Dodgson, J. Dickmann, A. Chessel, J. Dumais, and R.E.C. Salas. 2015. Wall mechanics and exocytosis define the shape of growth domains in fission yeast. Nat. Commun. 6:8400.

48. Davì, V., H. Tanimoto, D. Ershov, A. Haupt, H. De Belly, R. Le Borgne, E. Couturier, A. Boudaoud, and N. Minc. 2018. Mechanosensation Dynamically Coordinates Polar Growth and Cell Wall Assembly to Promote Cell Survival. Dev. Cell. 45:170–182.e7.

49. Odermatt, P.D., T.P. Miettinen, J. Lemière, J.H. Kang, E. Bostan, S.R. Manalis, K.C. Huang, and F. Chang. 2021. Variations of intracellular density during the cell cycle arise from tip-growth regulation in fission yeast. Elife. 10.

50. Weiss, J.N., and P. Korge. 2001. The cytoplasm: no longer a well-mixed bag. Circ. Res. 89:108–110.

51. Konopka, C.A., and S.Y. Bednarek. 2008. Variable-angle epifluorescence microscopy: a new way to look at protein dynamics in the plant cell cortex. Plant J. 53:186–196.

52. Weber, S.C., M.A. Thompson, W.E. Moerner, A.J. Spakowitz, and J.A. Theriot. 2012. Analytical tools to distinguish the effects of localization error, confinement, and medium elasticity on the velocity autocorrelation function. Biophys. J. 102:2443–2450.

53. Guigas, G., C. Kalla, and M. Weiss. 2007. Probing the nanoscale viscoelasticity of intracellular fluids in living cells. Biophys. J. 93:316–323.

54. Tolić-Nørrelykke, I.M., E.-L. Munteanu, G. Thon, L. Oddershede, and K. Berg-Sørensen. 2004. Anomalous diffusion in living yeast cells. Phys. Rev. Lett. 93:078102.

55. Weber, S.C., J.A. Theriot, and A.J. Spakowitz. 2010. Subdiffusive motion of a polymer *composed of subdiffusive monomers*. Phys. Rev. E Stat. Nonlin. Soft Matter Phys. 82:011913.

56. Ślęzak, J., and S. Burov. 2021. From diffusion in compartmentalized media to non-Gaussian random walks. Sci. Rep. 11:5101.

57. Santos, M.A.F. dos, M.A.F. dos Santos, and L.M. Junior. 2020. Log-Normal-Superstatistics for Brownian Particles in a Heterogeneous Environment. Physics. 2:571–586.

58. Bauer, D., H. Ishikawa, K.A. Wemmer, N.L. Hendel, J. Kondev, and W.F. Marshall. 2021. Analysis of biological noise in the flagellar length control system. iScience.24:102354.

59. Einstein, A. 1905. Über die von der molekularkinetischen Theorie der Wärme geforderte-Bewegung von in ruhenden Flüssigkeiten suspendierten Teilchen. Ann. Phys. 322:549–560.

60. Levene, H. 1960. Contributions to probability and statistics. Essays in honor of Harold-Hotelling. 278–292.

61. Weron, A., K. Burnecki, E.J. Akin, L. Solé, M. Balcerek, M.M. Tamkun, and D. Krapf. 2017. Ergodicity breaking on the neuronal surface emerges from random switching-between diffusive states. Sci. Rep. 7:5404.

62. Janczura, J., and A. Weron. 2015. Ergodicity testing for anomalous diffusion: small-sample statistics. J. Chem. Phys. 142:144103.

63. Mitchison, J.M. 1957. The growth of single cells: I. Schizosaccharomyces pombe. Exp.Cell Res. 13:244–262.

64. Mann, H., and D. Whitney. 1947. Controlling the false discovery rate: A practical and powerful approach to multiple testing. Ann. Math. Stat. 18:50–60.

65. Potma, E.O., W.P. de Boeij, L. Bosgraaf, J. Roelofs, P.J. van Haastert, and D.A. Wiersma. 2001. Reduced protein diffusion rate by cytoskeleton in vegetative and polarized dictyostelium cells. Biophys. J. 81:2010–2019.

66. Moeendarbary, E., L. Valon, M. Fritzsche, A.R. Harris, D.A. Moulding, A.J. Thrasher, E. Stride, L. Mahadevan, and G.T. Charras. 2013. The cytoplasm of living cells behaves as a poroelastic material. Nat. Mater. 12:253–261.

67. Brangwynne, C.P., G.H. Koenderink, F.C. MacKintosh, and D.A. Weitz. 2008. Cytoplasmic diffusion: molecular motors mix it up. J. Cell Biol. 183:583–587.

68. Fletcher, D.A., and R.D. Mullins. 2010. Cell mechanics and the cytoskeleton. Nature.463:485–492.

69. Weber, S.C., A.J. Spakowitz, and J.A. Theriot. 2012. Nonthermal ATP-dependent *fluctuations contribute to the in vivo motion of chromosomal loci*. Proc. Natl. Acad. Sci.U. S. A. 109:7338–7343.

70. Persson, L.B., V.S. Ambati, and O. Brandman. 2020. Cellular Control of Viscosity Counters Changes in Temperature and Energy Availability. Cell. 183:1572–1585.e16.

71. Okabe, K., N. Inada, C. Gota, Y. Harada, T. Funatsu, and S. Uchiyama. 2012. Intracellular temperature mapping with a fluorescent polymeric thermometer and fluorescence lifetime imaging microscopy. Nat. Commun. 3:705.

72. Hayashi, T., N. Fukuda, S. Uchiyama, and N. Inada. 2015. A cell-permeable fluorescent polymeric thermometer for intracellular temperature mapping in mammalian cell lines. PLoS One. 10:e0117677.

73. Chrétien, D., P. Bénit, H.-H. Ha, S. Keipert, R. El-Khoury, Y.-T. Chang, M. Jastroch, H.T. Jacobs, P. Rustin, and M. Rak. 2018. Mitochondria are physiologically maintained at close to 50 °C. PLoS Biol. 16:e2003992.

74. Wang, K., X.H. Sun, Y. Zhang, T. Zhang, Y. Zheng, Y.C. Wei, P. Zhao, D.Y. Chen, H.A. Wu, W.H. Wang, R. Long, J.B. Wang, and J. Chen. 2019. Characterization of cytoplasmic viscosity of hundreds of single tumour cells based on micropipette aspiration. R Soc Open Sci. 6:181707.

75. Yan, R., K. Chen, K. Xu, W. Li, and L. Xiang. 2020. Single-molecule displacement *mapping unveils nanoscale heterogeneities in intracellular diffusivity*. Nature Methods.17:524–530.

76. Gu, W., L.D. Etkin, M.A. Le Gros, and C.A. Larabell. 2007. X-ray tomography of Schizosaccharomyces pombe. Differentiation. 75:529–535.

77. Parkinson, D.Y., G. McDermott, L.D. Etkin, M.A. Le Gros, and C.A. Larabell. 2008. Quantitative 3-D imaging of eukaryotic cells using soft X-ray tomography. J. Struct. Biol.162:380–386.

78. Chaubet, L., A.R. Chaudhary, H.K. Heris, A.J. Ehrlicher, and A.G. Hendricks. 2020. Dynamic actin cross-linking governs the cytoplasm’s transition to fluid-like behavior. Mol. Biol. Cell. 31:1744–1752.

79. Schmitt, D.L., and S. An. 2017. Spatial Organization of Metabolic Enzyme Complexes in Cells. Biochemistry. 56:3184–3196.

80. Heinig, U., M. Gutensohn, N. Dudareva, and A. Aharoni. 2013. The challenges of *cellular compartmentalization in plant metabolic engineering*. Curr. Opin. Biotechnol.24:239–246.

81. Luby-Phelps, K., P.E. Castle, D.L. Taylor, and F. Lanni. 1987. Hindered diffusion of inert *tracer particles in the cytoplasm of mouse 3T3 cells*. Proc. Natl. Acad. Sci. U. S. A. 84:4910–4913.

82. Luby-Phelps, K., D.L. Taylor, and F. Lanni. 1986. Probing the structure of cytoplasm. J.Cell Biol. 102:2015–2022.

83. Arrio-Dupont, M., S. Cribier, G. Foucault, P.F. Devaux, and A. d’Albis. 1996. Diffusion of fluorescently labeled macromolecules in cultured muscle cells. Biophys. J. 70:2327–2332.

84. Arrio-Dupont, M., G. Foucault, M. Vacher, P.F. Devaux, and S. Cribier. 2000. Translational diffusion of globular proteins in the cytoplasm of cultured muscle cells. Biophys. J. 78:901–907.

85. Verkman, A.S. 2002. Solute and macromolecule diffusion in cellular aqueous compartments. Trends Biochem. Sci. 27:27–33.

86. Banks, D.S., and C. Fradin. 2005. Anomalous Diffusion of Proteins Due to Molecular Crowding. Biophysical Journal. 89:2960–2971.

87. Yu, I., T. Mori, T. Ando, R. Harada, J. Jung, Y. Sugita, and M. Feig. 2016. Biomolecular interactions modulate macromolecular structure and dynamics in atomistic model of a bacterial cytoplasm. Elife. 5.

88. Huang, J., Y. Huang, H. Yu, D. Subramanian, A. Padmanabhan, R. Thadani, Y. Tao, X. Tang, R. Wedlich-Soldner, and M.K. Balasubramanian. 2012. Nonmedially assembled *F-actin cables incorporate into the actomyosin ring in fission yeast*. J. Cell Biol.199:831–847.

89. Moreno, S., A. Klar, and P. Nurse. 1991. [56] Molecular genetic analysis of fission yeast Schizosaccharomyces pombe. In: Methods in Enzymology. Academic Press. pp. 795–823.

90. Maundrell, K. 1990. nmt1 of fission yeast. A highly transcribed gene completely repressed by thiamine. J. Biol. Chem. 265:10857–10864.

91. Sbalzarini, I.F., and P. Koumoutsakos. 2005. Feature point tracking and trajectory analysis for video imaging in cell biology. J. Struct. Biol. 151:182–195.

92. Schindelin, J., I. Arganda-Carreras, E. Frise, V. Kaynig, M. Longair, T. Pietzsch, S. Preibisch, C. Rueden, S. Saalfeld, B. Schmid, J.-Y. Tinevez, D.J. White, V. Hartenstein, K. Eliceiri, P. Tomancak, and A. Cardona. 2012. Fiji: an open-source platform for biological-image analysis. Nat. Methods. 9:676–682.

